# The unhindered lateral mobility of a rhomboid intramembrane protease enables synapse remodelling

**DOI:** 10.64898/2026.09.10.750656

**Authors:** Anna Bodzęta, Adam G. Grieve

## Abstract

Synapses must maintain their stable molecular organisation while retaining the capacity to remodel in response to changing demands. In order to flexibly change synaptic strength, the abundance of trans-synaptic adhesion complexes needs to be modulated, yet how this is achieved is poorly understood. Here, we identify the rhomboid protease RHBDL2 as a membrane-immersed regulator of adhesion complexes that enables mature synapses to adjust their composition both at steady-state and during their active restructuring. To do so, RHBDL2 possesses unrestrained diffusion properties that enable it to constitutively scan the neuronal surface and remove available adhesion molecules through irreversible cleavage of their transmembrane domains. Demonstrating its physiological importance, diffusion-dependent membrane surveillance by RHBDL2 is required for synapses to undergo remodelling during long term depression. Overall, we reveal a mechanism that cells use to preserve functional signalling complexes in membranes while selectively removing dispensable transmembrane proteins.

## Introduction

Effective and tunable transmission of information between neurons requires synapses to balance two seemingly opposing demands: preserving stable functional architecture while retaining the capacity for activity-dependent restructuring. Much of our understanding of synaptic regulation is derived from the study of postsynaptic neurotransmitter receptors and scaffolds via reversible post-translation modifications that coordinate their trafficking, exocytosis, endocytosis and autophagy^1–6^. However, changes in synaptic strength are accompanied by remodeling of transsynaptic adhesion complexes^7,8^, and considerably less is known about how this is regulated, or more broadly how persistent changes in synaptic architecture are established and maintained.

Synaptic adhesion molecules (CAMs) are particularly well positioned to coordinate this balance between stability and flexibility^9,10^. At mature synapses, CAMs serve both structural and signaling functions. Through interactions with intracellular scaffold proteins, they influence synaptic composition, including the number of glutamate receptors; through downstream signalling to the actin cytoskeleton, they shape dendritic spine morphology; and through transsynaptic interactions, they align neurotransmitter release sites with postsynaptic receptors^7–9,11^. A major unresolved question is how synaptic adhesion complexes are remodelled during different forms of synaptic restructuring. In particular, how can adhesion complexes be disassembled in a manner that not only disrupts existing interactions, but also prevents their immediate reassembly through either trans-and cis-interactions? More broadly, what mechanisms allow selected components of adhesion architecture to be durably removed while preserving its overall integrity and capacity for intercellular signalling?

We hypothesised that intramembrane proteolysis offers a mechanistic solution for this problem. Intramembrane proteases are evolutionarily conserved enzymes that irreversibly cleave transmembrane domains (TMDs) within the plane of the lipid bilayer^12,13^, providing a means of directly editing the membrane proteome. Its biological importance is underscored by the roles of the intramembrane proteases γ-secretase and SPPL2b in neurodegenerative conditions, such as Alzheimer’s disease^14–17^. In contrast, rhomboids (RHBDLs) are a distinct family of intramembrane protease that have not been studied in the context of mammalian neurons or synaptic transmission^18,19^. RHBDLs typically cleave type I TMDs through a catalytic serine-histidine dyad, and recognise substrates primarily through structural and conformational features of their TMD helices rather than short linear sequence motifs^20–22^. Their potentially broad spectrum of substrates that includes adhesion molecules^23^ known for their roles at synapses positions rhomboids as attractive candidates for the lateral regulation of synaptic architecture and function. This mechanism may be particularly well-suited to the neuronal plasma membrane, where specialised and crowded compartments such as the postsynaptic density are spatially segregated from defined zones for exocytosis and endocytosis^24–29^. Cleavage of synaptic CAMs within the plane of the plasma membrane could therefore selectively edit the local synaptic membrane proteome without requiring the removal through conventional trafficking pathways.

Here, using complementary imaging, biochemistry and biophysical approaches, we uncover a previously unrecognised mechanism for synaptic proteostasis mediated by the rhomboid protease RHBDL2. We show that RHBDL2 acts as a membrane-immersed editor of synaptic adhesion complexes, enabling synapse remodelling both at steady state and in response to plasticity-inducing stimuli. We propose that RHBDL2 employs unique diffusion properties to continuously scan the neuronal surface, removing adhesion molecules that are not engaged in stable transsynaptic complexes while allowing established complexes to persist. Overall, we reveal diffusion-dependent membrane surveillance as a mechanism for controlling synaptic protein composition and turnover, which is required to convert transient synaptic signals into persistent changes in synaptic state.

## Results

### RHBDL2 has the potential to regulate the architecture of mature excitatory synapses

To explore whether rhomboid intramembrane proteases can control synapse composition and architecture, we first examined the expression of the plasma membrane targeted rhomboids RHBDL2 and RHBDL3 during maturation of primary neurons. Quantitative RT-PCR of rat cortical neurons collected at different days *in vitro* (DIV) revealed a pronounced and progressive increase in rhomboid levels across neuronal maturation, with their relative expression reaching their highest levels at DIV19 (Figures 1A and S1A). Their preferential expression at late developmental stages indicate that their primary function may be linked to regulation of the surface proteome at mature synapses, rather than early neuronal development. As the most marked increase in expression was observed for RHBDL2, we focused on investigating its role in adjusting synapse architecture.

**Figure 1.**
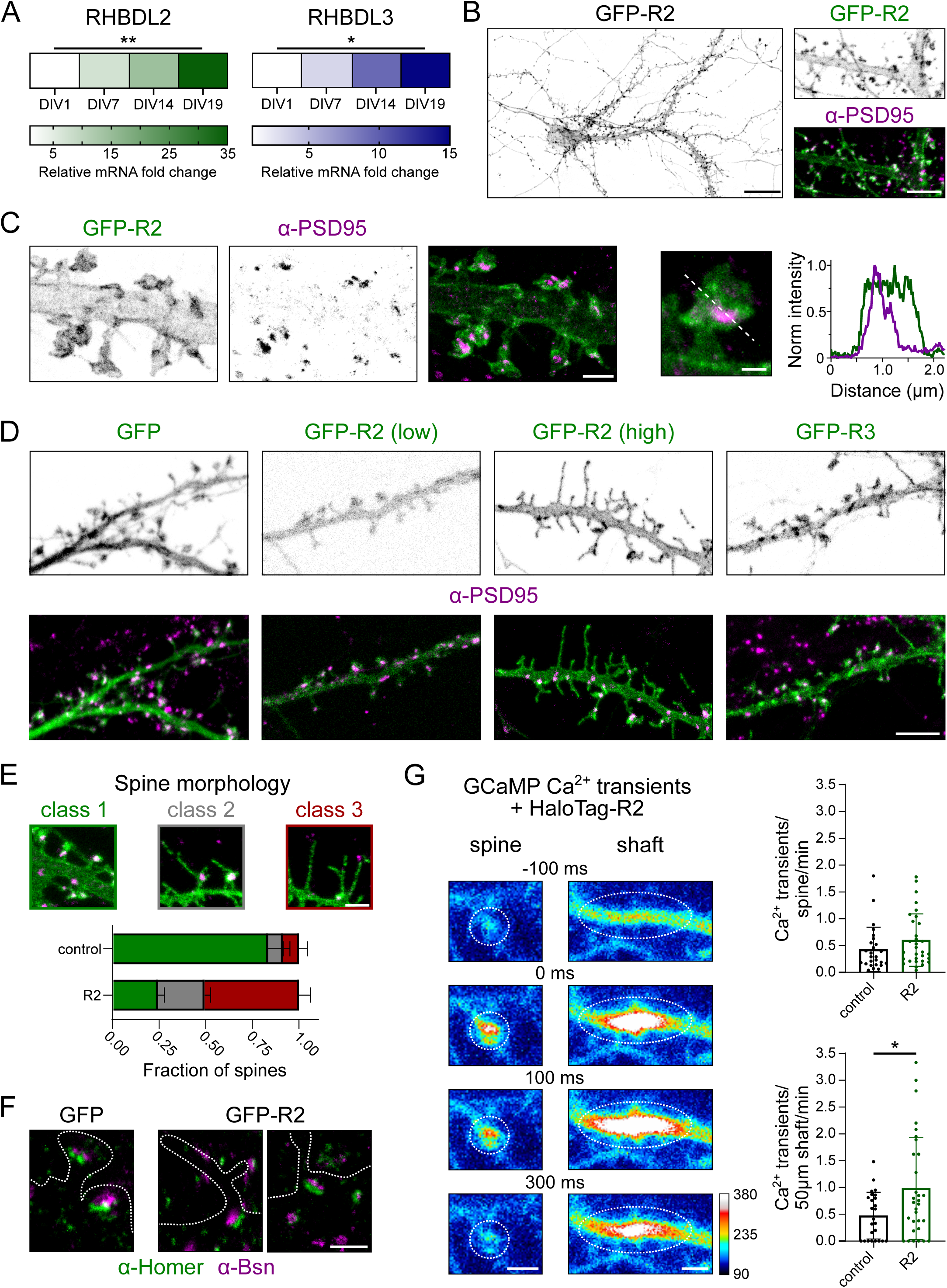
RHBDL2 has the potential to regulate the architecture of mature excitatory synapses. **(A)** Heat maps of relative fold changes of RHBDL2 and RHBDL3 mRNA expression in cortical neurons at different days in vitro (DIV). Mean values are included in Figure S1A. **(B)** Confocal image of hippocampal neurons expressing GFP-R2. Scale bar: 10 µm. Zoom image of GFP-R2 co-stained α-PSD95-S635P showing homogeneous distribution of R2 at dendritic plasma membrane. Scale bar: 5 µm. **(C)** Example gSTED image of a neuron expressing GFP-R2-S580 co-stained with postsynaptic marker α-PSD95-S635P. Scale bar: 2 µm. Example image and intensity line profile of individual dendritic spine. Scale bar: 500 nm. **(D)** Example confocal images of neurons expressing GFP, low or high levels of GFP-R2, or GFP-R3 co-stained with α-PSD95-S635P. Scale bar: 5 µm. **(E)** Classification of spine morphology at DIV19 in neurons transfected with GFP or GFP-R2 at DIV9. Example classes of spines in neurons expressing GFP-R2 and co-stained with α-PSD95-S635P. Class 1 – spine with a distinct PSD-containing head. Class 2 – spine with head with PSD and spinules. Class 3 – filopodia-like spine without a PSD. Scale bar: 2 µm. Mean fraction values ± SD (a.u.): ctrl (GFP) class 1 0.84 ± 0.08, class 2 0.08 ± 0.04; class 3 0.09 ± 0.05; GFP-R2 class 1 0.24 ± 0.03, class 2 0.25 ± 0.03, class 3 0.51 ± 0.06. N = 3 independent experiments. **(F)** Examples of gSTED images of neurons expressing GFP or high levels of GFP-R2 co-stained with postsynaptic marker α-Homer1-S580 and presynaptic marker α-Bassoon-S635P (Bsn). Dashed lines represent outlines of GFP-positive neurons. Scale bar: 1 µm. **(G)** Example images from GCaMP6f time series experiments in neurons expressing HaloTag-R2 showing Ca^2+^ events at spines and shafts. HaloTag-R2 was labeled with ligand JFX650. Royal LUT was used to display changes in fluorescence intensity. Dotted circles represent Ca^2+^ events. Scale bar: 2 µm. Quantification of Ca^2+^ transient frequency in control neurons and neurons expressing HaloTag-R2. Mean frequency values ± SD: spine (transient/spine/min): ctrl 0.43 ± 0.41, R2 0.6 ±0.49; shaft (transient/50 µm/min): ctrl 0.48 ± 0.44, R2 0.98 ± 0.95. N = 6 independent experiments. Unpaired t-test followed by Welch’s correction.

The spatial distribution of proteins within the highly organised neuronal plasma membrane ultimately determines their function, therefore, we studied the subcellular localisation of RHBDL2 in rat hippocampal neurons. GFP-tagged RHBDL2 was broadly distributed within the neuronal plasma membrane, including axons, dendrites and dendritic spines, without apparent enrichment within any specific membrane compartment (Figure 1B). Knock-down of endogenous RHBDL2 and re-expression of shRNA-sensitive GFP-RHBDL2 confirmed that this broad distribution was not an artefact of ectopic expression (Figure S1B-D). Next, to resolve whether RHBDL2 was also homogenously distributed across synaptic membrane compartments, its localisation was assessed at nanoscale resolution with two-color gated stimulated emission depletion (gSTED) microscopy, using PSD95 as a postsynaptic density marker. RHBDL2 was consistently detected within synaptic spines and overlapped with PSD95-containing synapses. RHBDL2 was not concentrated within the PSD itself but was uniformly distributed across the membranes of the dendritic spine and shaft (Figure 1C). This broad localisation indicates that RHBDL2 has unusual access to both synaptic and extrasynaptic substrates of the neuronal surface.

Interestingly, we observed striking heterogeneity in dendritic spine morphology, which was dependent on the level of GFP-RHBDL2 expression. Neurons expressing higher levels of RHBDL2 exhibited a marked increase in elongated filopodia-like protrusions, with a corresponding reduction in mature mushroom-like spines (Figure 1D and 1E). This structural remodelling was accompanied by a redistribution of the postsynaptic marker PSD95 from discrete puncta in dendritic spines to the dendritic shaft. This suggested that RHBDL2 not only influences spine morphology, but also the positioning of excitatory synapses.

To verify whether these shaft-associated PSD95 puncta represented functional synapses, we labelled neurons for an alternative postsynaptic marker, Homer1, together with the presynaptic active zone marker Bassoon, and performed gSTED imaging. The majority of Homer1-positive puncta in the dendritic shaft were closely apposed to Bassoon-positive presynaptic terminals (Figure 1F), indicating that RHBDL2 expression does not simply disperse postsynaptic proteins or wholesale uncouple synapses, but it actively reshapes spine structure and synapse positioning.

To test whether these changes in synapse position and spine morphology influence synaptic signaling, we measured spontaneous postsynaptic Ca^2+^ transients using the fluorescence intensity-based sensor GCaMP6f^30^. In neurons with elevated levels of RHBDL2, the frequency of Ca^2+^ events at intact spines was not altered, however we observed an increased number of Ca^2+^ transients in the dendritic shaft that displayed distinct and prolonged kinetics to those at spines (Figure 1G and S1F). This demonstrated that RHBDL2-elicited shaft synapses are functional, but the spatial constraint of their downstream calcium signalling processes is lost.

Notably, expression of the neuronal rhomboid protease RHBDL3 failed to alter spine morphology and synapse position (Figure 1D), indicating that this phenotype is not a general consequence of increased rhomboid protease activity, but instead reflects a specific function of RHBDL2.

We next explored whether this phenotype reflected an effect on spine morphogenesis or whether RHBDL2 could remodel established synapses. To distinguish between these possibilities, neurons were transfected either before the onset of synaptogenesis (DIV9) or after mature synapse had formed (DIV16)^31^, and spine morphology was analysed at DIV19. Elevated RHBDL2 expression increased the proportion of filopodia-like protrusions lacking postsynaptic markers under both conditions (Figures 1E and S1E). This demonstrated that RHBDL2 can reshape pre-existing mature spines.

Together, these findings identify RHBDL2 as a neuronal plasma membrane protease with the innate ability to regulate the structural organisation of mature synapses. Its uniform distribution along the neuronal surface and at synapses, together with its ability to remodel spine architecture, suggest RHBDL2 acts on membrane proteins that control synaptic stability.

### RHBDL2 remodels synapses by proteolytically editing synaptic CAMs

The pronounced remodelling of spine morphology and synapse position by RHBDL2 suggested that the intramembrane protease acts on membrane proteins that stabilise mature synapses. As described earlier, synaptic CAMs are attractive candidates due to their co-ordinatation of trans-synaptic adhesion, and regulation of synaptic maturation and maintenance^8,9,11^ (Figure 2A). Furthermore, their disruption results in elongated, irregularly shaped protrusions^32–34^ resembling those observed following RHBDL2 expression. As adhesion proteins are widespread and evolutionarily conserved rhomboid substrates^35–37^, we hypothesised that RHBDL2 remodels synapses by proteolytically regulating the abundance and localisation of synaptic CAMs.

**Figure 2.**
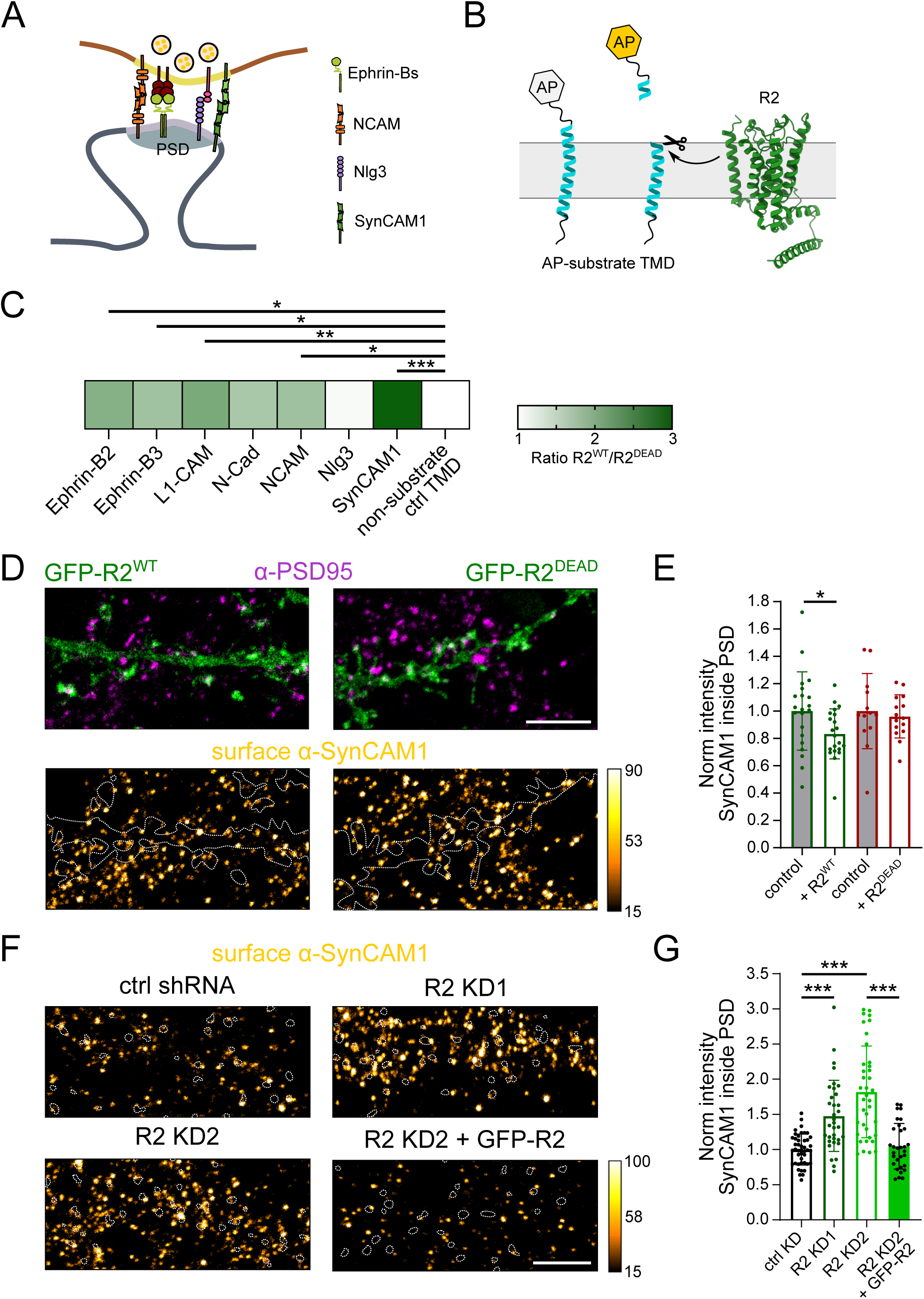
RHBDL2 proteolytically edits synaptic CAMs **(A)** Schematic of selected synaptic CAMs. **(B)** Schematic illustration of the AP-TMD shedding assay. **(C)** Heat map showing relative cleavage of selected CAM TMDs by R2 in HEK293T cells. Mean values of R2^WT^/R2^DEAD^ ratio ± SD (a.u.) for tested TMDs: ephrin-B2 1.95 ± 0.36, ephrin-B3 1.73 ± 0.51, L1-CAM 2.04 ± 0.53, N-Cad 1.67 ±0.38, NCAM 1.74 ± 0.36, Nlg3 1.06 ± 0.18, SynCAM1 2.92 ± 0.42, non-substrate TMD 0.85 ± 0.29. N = 3 or 4 independent experiments. One-way ANOVA followed by Dunnett’s multiple comparisons test. **(D)** Example confocal images of neurons expressing GFP-R2^WT^ or GFP-R2^DEAD^ co-stained with PSD marker α-PSD95-S580 and corresponding images of surface α-SynCAM1-STAR-RED staining. Orange Hot LUT was used for SynCAM1 images to display differences in fluorescence intensity. Dashed lines represent outlines of GFP positive neurons. Scale bar: 5 µm. Calibration bar: grey scale values. **(E)** Quantification of the intensity of surface α-SynCAM1 inside the PSD in neurons expressing GFP-R2^WT^ or GFP-R2^DEAD^ normalised to the intensity inside the PSD of non-transfected neurons from the same field of view. Mean intensity values ± SD (a.u.): ctrl 1 ± 0.25, + R2^WT^ 0.82 ± 0.18; ctrl 1 ± 0.24, + R2^DEAD^ 0.97 ± 0.17. N = 4 independent experiments. One-way ANOVA followed by Sidak’s multiple comparisons test. **(F)** Example confocal images of surface α-SynCAM1-STAR-RED staining in neurons expressing control shRNA, neurons with R2 KD1, KD2 and R2 KD2 with re-expression of GFP-R2. Orange Hot LUT was used to display differences in fluorescence intensity. Dotted circles represent outlines of PSDs based on corresponding images of α-PSD95-S580. Scale bar: 5 µm. **(G)** Quantification of the intensity of surface α-SynCAM1 inside the PSD of neurons transduced with R2 KD1, KD2 and R2 KD2 with re-expression of GFP-R2 normalised to intensity inside the PSD of neurons expressing control shRNA. Mean intensity values ± SD (a.u.): ctrl KD 1 ± 0.22, R2 KD1 1.48 ± 0.5, R2 KD2 1.82 ± 0.65, R2 KD2 + GFP-R2 1.05 ± 0.32. N = 4 independent experiments. One-way ANOVA followed by Sidak’s multiple comparisons test.

To initially test this hypothesis, we performed a cell biological substrate screen using transmembrane domains (TMDs) of representative CAMs from multiple families in cell lines. Alkaline phosphatase (AP) was fused to the extracellular domain of the TMDs of ephrin-B2, ephrin-B3, L1-CAM, N-cadherin (N-Cad), NCAM, Neuroligin-3 (Nlg3) and SynCAM1, together with the non-substrate control TMD of Calnexin. AP-TMD constructs were co-expressed with either active or catalytically dead RHBDL2 and intramembrane cleavage was quantified using an established AP release assay^21^ (Figure 2B). RHBDL2 efficiently cleaved five of the seven CAMs tested, including ephrin-B2, ephrin-B3, L1-CAM, NCAM, SynCAM1, whereas cleavage of Nlg3 and N-Cad was not significantly above a non-substrate control, Calnexin (Figure 2C). These findings demonstrate that RHBDL2 possesses broad substrate specificity toward the TMDs of multiple classes of synaptic adhesion molecules, suggesting it may regulate the composition of the neuronal surface through coordinated proteolytic turnover of adhesion proteins.

Among the identified substrate TMDs, SynCAM1 exhibited the highest level of RHBDL2-dependent cleavage. Given the well-established role of SynCAM1 in stabilising mature excitatory synapses^32,38–41^, we selected this CAM for physiological validation in neurons. To determine whether SynCAM1 is a physiological substrate of RHBDL2 at the neuronal plasma membrane, we quantified endogenous surface levels of SynCAM1 in hippocampal neurons expressing either active or catalytically dead RHBDL2. Expression of active RHBDL2 significantly reduced the intensity of SynCAM1 within the PSD, whereas the inactive enzyme had no detectable effect (Figure 2D and 2E). Thus, RHBDL2 is able to proteolytically regulate surface levels of endogenous SynCAM1 at synapses.

We next asked whether endogenous RHBDL2 constitutively controls surface CAM abundance. We used two independent shRNAs to efficiently reduce RHBDL2 expression in cultured neurons (Figure S1B). Upon RHBDL2 depletion, we observed no discernable difference in spine morphology, synapse positioning nor in spontaneous calcium signaling (Figure S2A and S2B). But, as would be predicted for a rhomboid substate, RHBDL2 knock-down resulted in a significant accumulation of surface SynCAM1 at excitatory synapses compared with neurons expressing a control shRNA (Figure 2F and 2G). Importantly, re-expression of shRNA-sensitive GFP-RHBDL2 restored synaptic SynCAM1 intensity to that observed in control, demonstrating that this phenotype specifically results from loss of RHBDL2 function (Figure 2G).

Together, these findings identify SynCAM1 as a physiological synaptic substrate of RHBDL2, and indicate that RHBDL2 continually regulates the abundance of synaptic adhesion molecules through direct intramembrane proteolysis. More broadly, the ability of RHBDL2 to cleave the TMDs of multiple synaptic CAM families suggests that it functions as a proteolytic editor of synapse architecture, rather than a regulator of any specific adhesion protein or family. Such control of synaptic adhesion would be expected to promote remodelling of mature synapses during activity-dependent restructuring, which led us to ask whether RHBDL2 is required for synaptic plasticity.

### RHBDL2 is required to maintain the structural flexibility necessary for synaptic remodelling

Because dynamic changes in synaptic adhesion strength are an essential feature of activity-dependent plasticity^7,42^, we hypothesised that RHBDL2 maintains the remodelling capacity of mature synapses. To test this idea, we examined whether RHBDL2 is required for changes in synapse composition during LTD, where persistent reductions in synaptic transmission are accompanied by structural remodelling^2,42^. To this end, we used the removal of synaptic AMPA receptors (AMPARs) during metabotropic glutamate receptor (mGluR)-dependent LTD as a proxy^43^. Chemical mGluR-LTD (cLTD) was induced in cultured hippocampal neurons by treatment with the group I mGluR agonist S-(3,5)-DHPG, and levels of surface AMPARs within the PSD were quantified, following shRNA-mediated depletion (Figure 3A). As expected, neurons expressing a control shRNA exhibited a robust reduction in synaptic surface AMPARs following DHPG treatment, consistent with efficient induction of cLTD. In contrast, depletion of RHBDL2 largely abolished this response (Figure 3B and 3C). Following cLTD induction, synaptic AMPAR levels remained comparable to untreated neurons, indicating that LTD-triggered remodelling of excitatory synapses fails to occur in the absence of RHBDL2.

**Figure 3.**
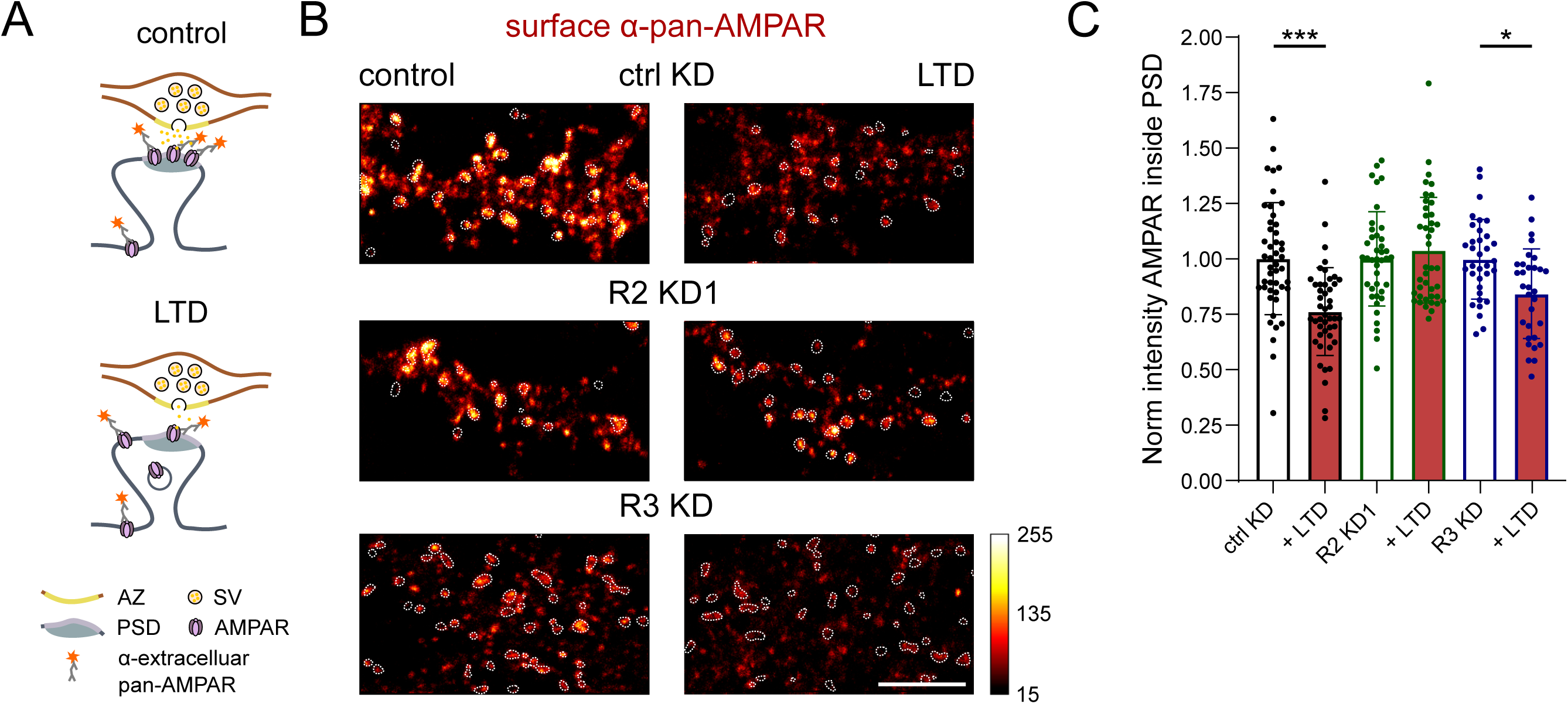
RHBDL2 is required for synapse remodeling during LTD. **(A)** Schematic illustration of surface labeling of AMPAR during LTD. **(B)** Example images of surface α-pan-AMPAR-S580 in control conditions and after LTD induction in control neurons, or neurons with R2 KD or R3 KD. Red Hot LUT was used to display differences in fluorescence intensity. Dotted circles represent outlines of PSDs based on corresponding images of α-PSD95-S635P. Note that for all LTD experiments the nanobody α-PSD95-S635P was used. Scale bar: 5 µm. Calibration bar: grey scale values. **(C)** Quantification of surface α-pan-AMPAR inside the PSD after induction of LTD normalised to control levels. Mean intensity values ± SD (a.u.): ctrl KD 1.00 ± 0.27, cLTD ctrl KD 0.79 ± 0.20; ctrl R2 KD1 1.00 ± 0.21, cLTD R2 KD1 1.03 ± 0.24; ctrl R3 KD 1.00 ± 0.18, cLTD R3 KD 0.84 ± 0.20. N = 5 or 6 independent experiments. One-way ANOVA followed by Sidak’s multiple comparisons test.

To determine whether this phenotype was specific to RHBDL2, or whether it reflects a more general role for rhomboid proteases in neurons, we performed the same experiment following the knockdown of the neuron-specific rhomboid protease, RHBDL3. In contrast to RHBDL2 depletion, RHBDL3 loss had no detectable effect on AMPAR removal during cLTD (Figure 3B and 3C), confirming a specific role for RHBDL2 in this form of synaptic remodelling.

Overall, these findings establish RHBDL2 as a regulator of synapse stability. Rather than simply controlling the abundance of individual adhesion proteins, RHBDL2 appears to preserve the capacity of mature synapses to undergo dynamic remodelling. As RHBDL2 is constitutively primed for substrate engagement and does not require external signals for activity^21,44^, these observations support a model in which continuous proteolytic turnover of synaptic CAMs prevents excessive stabilisation of trans-synaptic adhesion complexes, thereby maintaining structural flexibility that is required for regulation of synaptic glutamate receptors during LTD.

### RHBDL2 exhibits exceptionally rapid diffusion in neuronal membranes

Synaptic restructuring is a highly dynamic event that occurs at defined plasma membrane locations. In order to regulate this process, RHBDL2 must have properties that allow it to laterally interrogate synaptic content for substrates. Previous studies in artificial membranes and non-neuronal cells showed that the rhomboid superfamily of proteins generally diffuse rapidly, which may explain its homogenous distribution at the neuronal plasma membrane (Figure 1B and 1C), but more generally its physiological importance has not been established^45^. We therefore examined the diffusion behaviour of RHBDL2 at high spatial and temporal resolution, using single-molecule tracking (SMT).

To do so initially, HaloTag-labelled RHBDL2 was benchmarked against other members of rhomboid superfamily: RHBDL1, RHBDL3, the pseudoprotease iRhom1, as well as the unrelated GxGD intramembrane protease SPPL2b in U2OS cells. Using a picomolar concentration of the HaloTag ligands JFX650 or JF635i, we achieved stochastic labeling and robust detection of single molecules of these proteins at the plasma membrane. Individual trajectories were reconstructed and used to calculate instantaneous diffusion coefficients (D_inst_) (Figure 4A). All imaged intramembrane proteases and pseudoproteases substantially differed in their diffusion behaviour at the plasma membrane. Notably, RHBDL2 displayed the highest D_inst_, approaching that of the freely diffusing single-pass transmembrane protein GT46 (Figure 4B and 4C). In contrast, RHBDL1, RHBDL3, iRhom1 and SPPL2b all exhibited significantly lower mobility (Figure 4B and 4C). Thus, rapid lateral diffusion is not a general feature of intramembrane proteases, but instead represents a distinctive property of RHBDL2.

**Figure 4.**
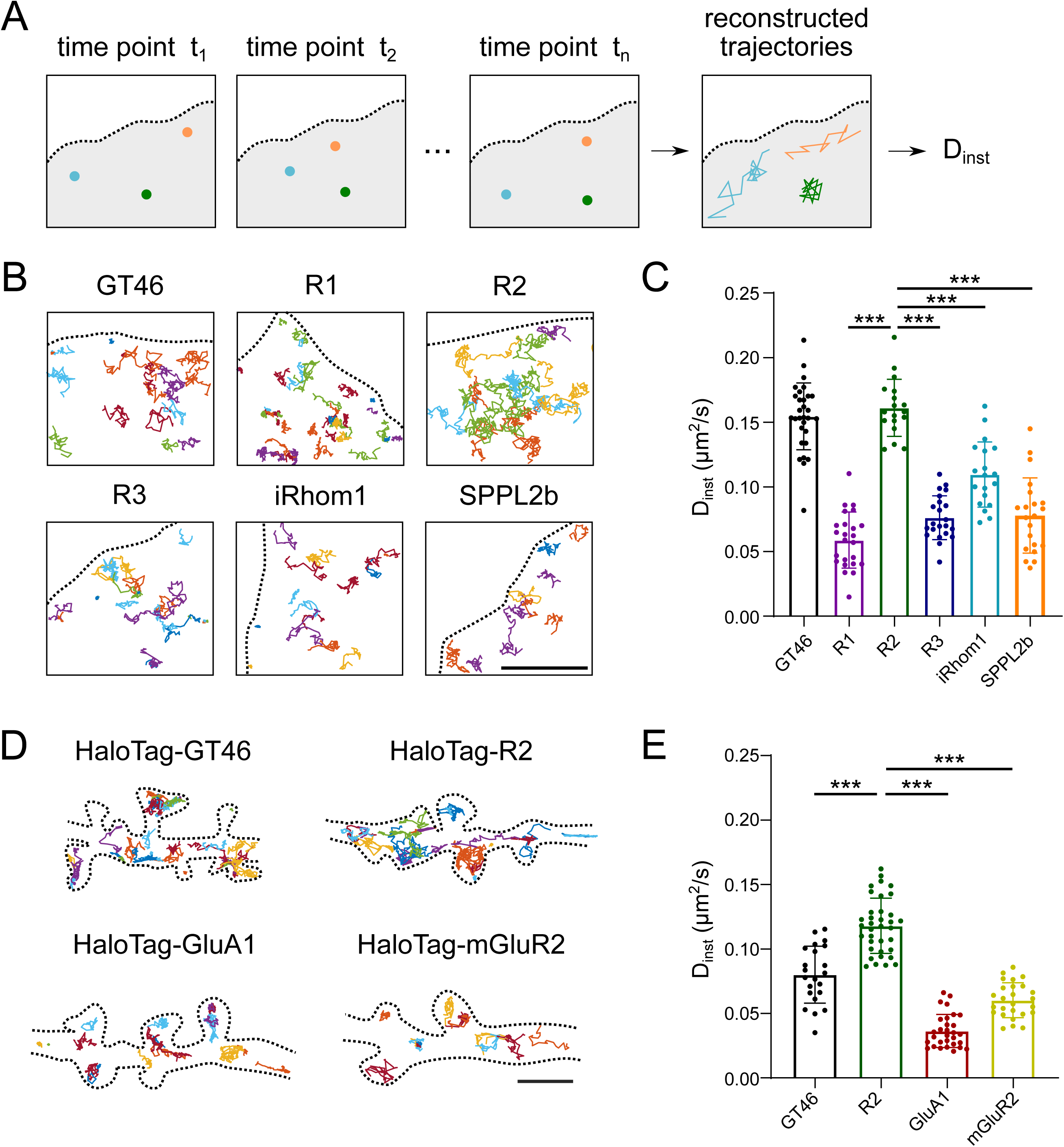
RHBDL2 exhibits exceptionally rapid diffusion in cellular membranes. **(A)** Schematic illustration of SMT experiments. **(B)** Example trajectories of HaloTag-tagged GT46, rhomboids RHBDL1 (R1), RHBDL2 (R2) and RHBDL3 (R3), pseudoprotease iRhom1, and GxGD protease SPPL2b in U2OS cells. Trajectories are displayed in random colors. Dashed lines represent the outline of U2OS cells. HaloTag ligand JFX650 was used for tracking of GT46, R1, R2, R3 and SPPL2b. Note that for tracking of iRhom1 cell impermeable HaloTag ligand JF635i was used to assess mobility of only the surface pool of this protein, due to its significant internal pool. Scale bar: 3 µm. **(C)** Quantification of average instantaneous D_inst_ of HaloTag-tagged proteins from **(B)**. Mean D_inst_ values ± SD (µm^2^/s): GT46 0.15 ± 0.03, R1 0.06 ± 0.02, R2 0.16 ± 0.02, R3 0.08 ± 0.02, iRhom1 0.11 ± 0.03, SPPL2b 0.08 ± 0.03. N = 3 independent experiments. One-way ANOVA followed by Dunnett’s multiple comparison test. **(D)** Example trajectories of HaloTag-tagged GT46, RHBDL2, GluA1 and mGluR2 in hippocampal neurons. Trajectories are displayed in random colors. Dashed lines represent outlines of transfected neurons. Note that for tracking of GluA1 and mGluR2 cell impermeable HaloTag ligand JF635i was used to assess mobility of only surface pool of these receptors, due to their significant internal pool. Scale bar: 2 µm. **(E)** Quantification of average instantaneous D_inst_ of HaloTag-tagged proteins from **(D)**. Mean D_inst_ values ± SD (µm^2^/s): GT46 0.08 ± 0.02, R2 0.12 ± 0.02, GluA1 0.04 ± 0.01, mGluR2 0.06 ± 0.01. N = 3 – 5 independent experiments. One-way ANOVA followed by Sidak’s multiple comparisons test.

We next asked whether this remarkably high diffusion rate is maintained within the crowded environment of the dendritic plasma membrane, which is itself compartmentalised into multiple specialised membrane domains^1,28^. Trajectory maps obtained from SMT in hippocampal neurons revealed that RHBDL2 diffuses freely throughout dendritic shafts and can enter and exit spines, without obvious confinement to specific membrane compartments (Figure 4D). This unimpeded diffusion was consistent with the homogenous distribution of RHBDL2 observed in fixed neurons by gSTED imaging (Figure 1C), and reveals that RHBDL2 continuously scans both synaptic and extrasynaptic regions of the neuronal surface. Strikingly, RHBDL2 displayed relatively higher mobility than single-pass GT46 in the neuronal plasma membrane^46^ (Figure 4D, 4E and Movie S1). RHBDL2 also diffused much more rapidly than other multi-pass transmembrane synaptic proteins, such as the GluA1 subunit of AMPARs, and the metabotropic glutamate receptor mGluR2 (Figure 4D and 4E). Altogether, these observations suggest that exceptional lateral mobility is an intrinsic property of RHBDL2 that is preserved, and potentially selected for, in the complex environment of the neuronal surface.

These data indicated that this unique diffusion behaviour is a biophysical property of the protease which may be essential for the neurobiological function of RHBDL2. We therefore next asked whether restricting RHBDL2 mobility alters its ability to regulate synaptic architecture.

### Rapid lateral RHBDL2 diffusion is important for shaping synapse architecture

To test directly whether the rapid lateral mobility of RHBDL2 is an important determinant of its function, we sought to immobilise RHBDL2 by anchoring it to the cortical actin cytoskeleton using the calponin-homology domain from utrophin^47^ (UR2, Figure 5A). Using SMT, we confirmed that UR2 displayed a markedly reduced D_inst_ and fraction of mobile trajectories, compared to the wild type enzyme (Figure 5B, 5C and Movie S2). This reduced mobility resulted from our intended actin-dependent confinement strategy, as brief disruption of the cortical actin cytoskeleton with a low concentration of latrunculin B led to significant release of this pool of immobilised RHBDL2 (Figure 5C). Thus, fusion to utrophin effectively restricted the spatial exploration of RHBDL2 without altering its localisation to the neuronal plasma membrane.

**Figure 5.**
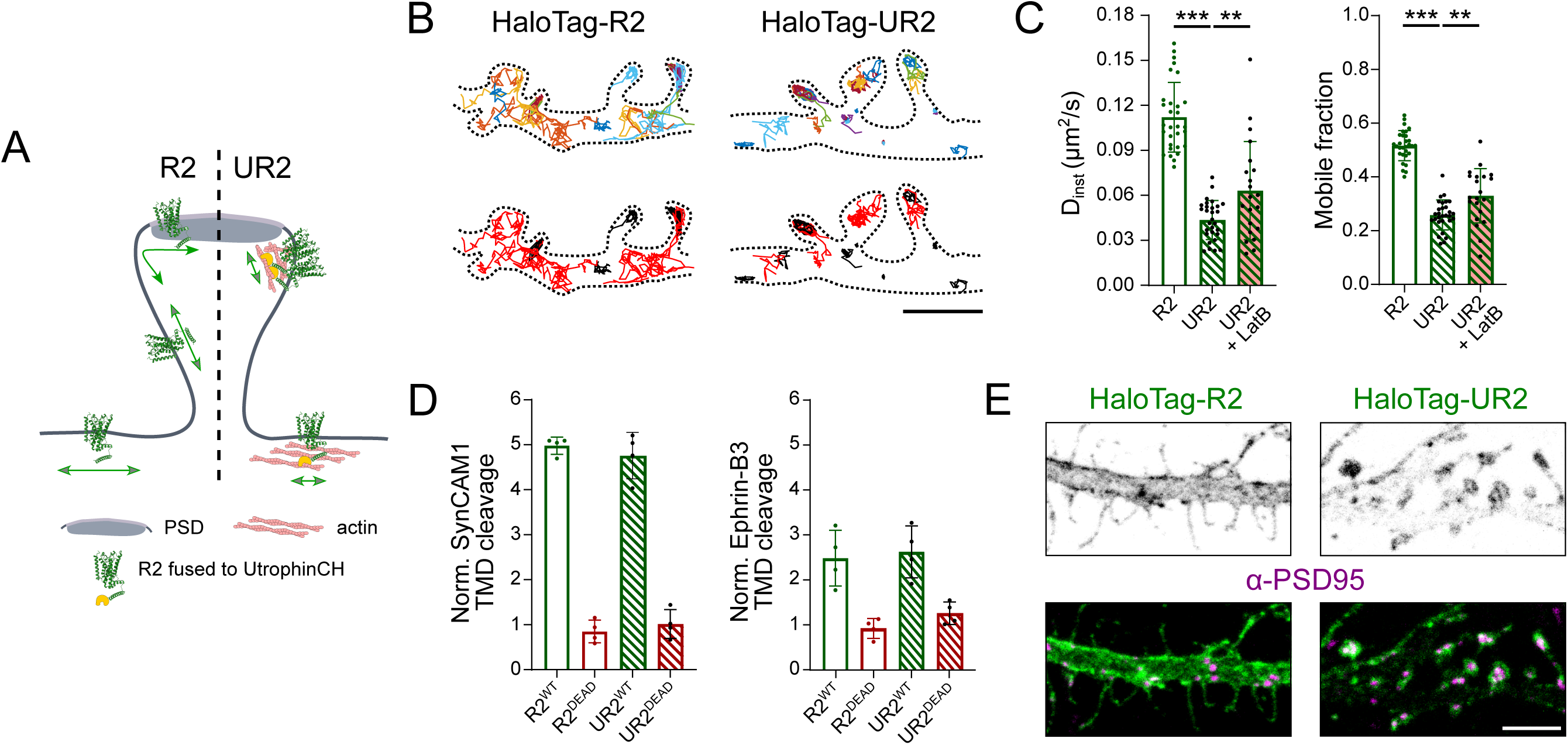
An immobile form of RHBDL2 retains activity but fails to reshape synaptic architecture. **(A)** Schematic of the intended effect of restricting the localisation and mobility of R2 through anchoring to the actin cytoskeleton via utrophinCH (UR2). **(B)** Example trajectories of HaloTag-tagged R2 and UR2 in hippocampal neurons; top: individual trajectories displayed in random colors; bottom: red for mobile trajectories, black for immobile trajectories. Dashed lines represent outlines of transfected neurons. HaloTag ligand JFX650 was used for SMT. Scale bar: 2 µm. **(C)** Quantification of the average instantaneous D_inst_ and mobile fraction of HaloTag-tagged R2, UR2 and UR2 after treatment with LatB (1 µM, 2-5 min). Mean D_inst_ values ± SD (µm^2^/s): D_inst_: R2 0.11 ± 0.02 µm^2^/s, UR2 0.04 ± 0.01 µm^2^/s, UR2 + LatB 0.06 ± 0.03 µm^2^/s. Mean mobile fraction ± SD (a.u.): R2 0.52 ± 0.06, UR2 0.26 ±0.06, UR2 + LatB 0.33 ± 0.10. N = 3 independent experiments. One-way ANOVA followed by Sidak’s multiple comparisons test. **(D)** Quantification of cleavage of SynCAM1 and ephrin-B3 TMDs by R2 or UR2 in HEK293T cells. Mean relative cleavage ± SD (a.u.): SynCAM1 TMD + R2^WT^ 5.00 ± 0.19, + R2^DEAD^ 0.85 ±0.25, + UR2^WT^ 4.8 ± 0.52, + UR2^DEAD^ 1.01 ±0.32; Ephrin-B3 TMD + R2^WT^ 2.48 ± 0.62, + R2^DEAD^ 0.92 ±0.22, + UR2^WT^ 2.62 ± 0.58, + UR2^DEAD^ 1.26 ±0.25. N = 4 independent experiments. One-way ANOVA followed by Sidak’s multiple comparisons test. **(E)** Example confocal images of neurons expressing HaloTag-R2 or HaloTag-UR2 co-stained with α-PSD95-S635P. Note that HaloTag was labeled using α-HaloTag antibody. Scale bar: 5 µm.

To determine whether reduced mobility affected RHBDL2 catalytic activity, we compared cleavage of SynCAM1 and ephrin-B3 TMDs using the AP shedding assay. Wild-type and immobilised RHBDL2 cleaved both substrates with indistinguishable efficiency (Figure 5D), demonstrating that immobilising RHBDL2 via tethering to the actin cytoskeleton does not impair its intrinsic enzymatic activity. This separation of catalytic activity from membrane mobility provided a unique opportunity to determine whether rapid diffusion itself contributes to RHBDL2 function.

To test our hypothesis that rapid lateral mobility of RHBDL2 is essential for its synaptic function, we examined whether diffusion is required for RHBDL2-dependent remodelling of dendritic spines and synapse positioning. Whereas expression of wild-type RHBDL2 reproduced the characteristic increase in elongated filopodia-like protrusions together with the redistribution of synapse to the dendritic shaft, expression of the immobilised form of RHBDL2, UR2, had no effect on the morphology of dendritic spines nor on synapse positioning (Figure 5E). Consistent with its interaction with cortical actin, immobilised RHBDL2 accumulated in actin-rich spine heads. Because utrophin expression has previously been reported to influence actin dynamics^48^, we questioned whether the absence of a synaptic morphology phenotype could result simply from stabilisation of the actin cytoskeleton in spines. Independent co-expression of GFP-utrophinCH with freely diffusing HaloTag-RHBDL2 did not prevent RHBDL2-induced spine remodelling (Figure S3), demonstrating that the loss of function observed with UR2 arises from restricted membrane mobility rather than secondary effects on spine actin organisation. Moreover, these data indicate that catalytic activity alone is insufficient to remodel synapses in the absence of rapid lateral diffusion.

### RHBDL2 maintains synaptic remodelling competence through lateral surveillance of adhesion molecules

Having established that membrane diffusion is required for RHBDL2-dependent structural remodelling of synapse, we next asked whether its mobility affects substrate cleavage in the complex endogenous membrane protein landscape of the neuronal surface.

SynCAM1 predominantly localises within the periphery of the PSD^41^, where it is engaged trans-synaptically with presynaptic CAMs to stabilise mature synapses^32,40^. We noted in our previous experiments that RHBDL2 depletion appeared to increase both the synaptic and wider dendritic pools of SynCAM1 (Figure 2F – R2 KD). This suggested that RHBDL2 regulates synaptic CAMs that are distributed across the entirety of the neuronal surface. We therefore hypothesised that rapid diffusion enables RHBDL2 to continuously encounter and eliminate non-synaptic pools of SynCAM1 molecules.

To test this idea, we quantified surface SynCAM1 more generally in the whole dendritic plasma membrane following expression of either wild-type or immobilised RHBDL2. As expected, expression of freely diffusing wild-type RHBDL2 significantly decreased SynCAM1 intensity throughout the neuronal cell surface, in a manner dependent on its catalytic activity. Interestingly, the cell surface-immobilised yet catalytically active form of RHBDL2, UR2, had no effect on surface SynCAM1, essentially phenocopying inactive RHBDL2 (Figure 6A and 6B). Although this confirmed our hypothesis, it was a somewhat paradoxical observation: immobile RHBDL2 failed to process SynCAM1 despite its efficient targeting to the plasma membrane. Mechanistic insight into the inability to process these non-synaptic pools of endogenous SynCAM1 was provided by gSTED imaging, which revealed that immobile RHBDL2 specifically accumulated around the postsynaptic density, and was excluded from the PSD itself (Figure 6C), in line with the known enrichment of actin in dendritic spines^49^. These findings confirm that restricting RHBDL2 diffusion limits the spatial range over which the protease can regulate substrate abundance.

**Figure 6.**
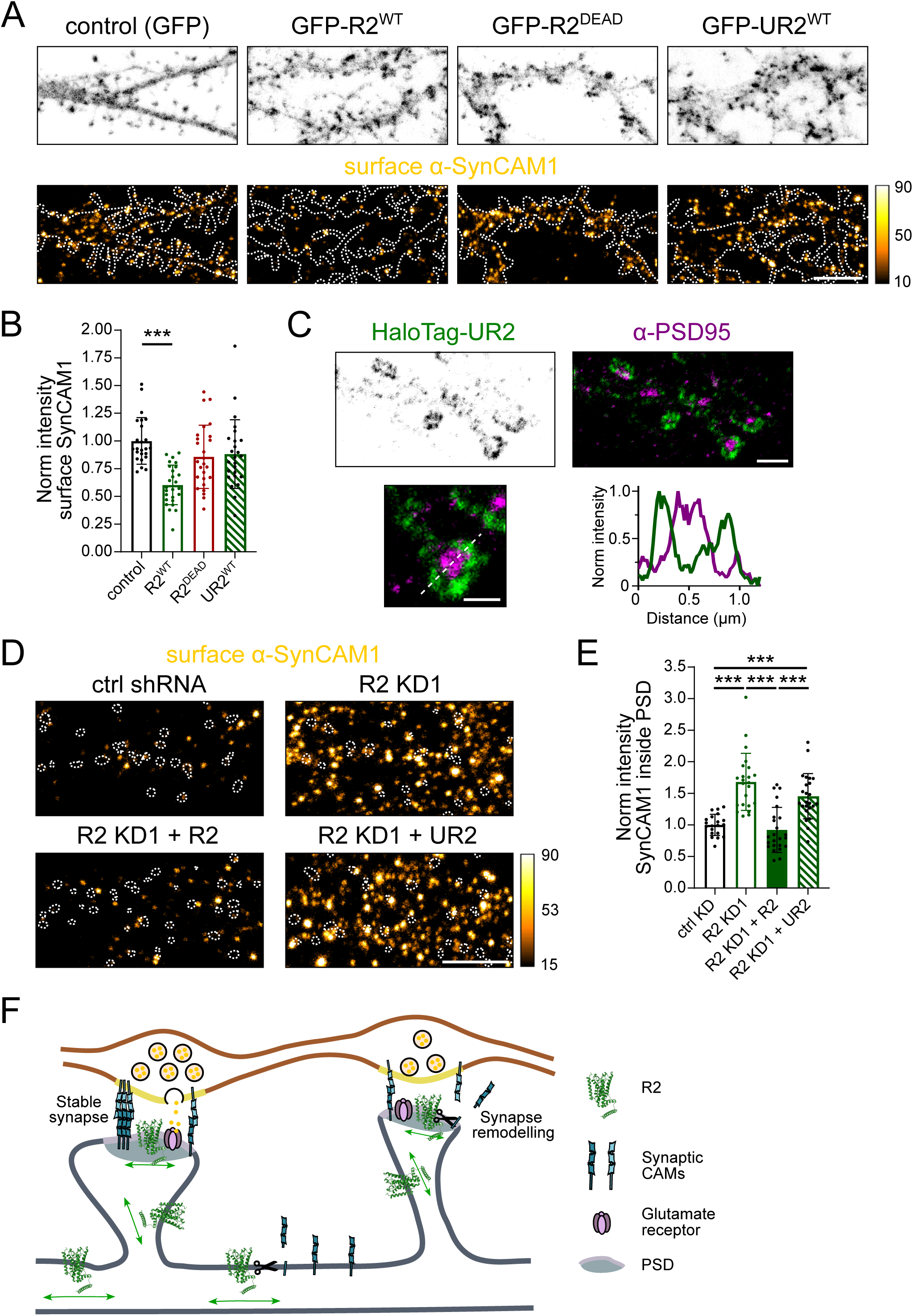
Rapid lateral diffusion of RHBDL2 determines its synaptic function. **(A)** Example confocal images of neurons expressing GFP, GFP-R2^WT^, GFP-R2^DEAD^ or GFP-UR2^WT^ and corresponding images of surface α-SynCAM1-STAR-RED staining. Orange Hot LUT was used for SynCAM1 images to display differences in fluorescence intensity. Dashed lines represent outlines of GFP-positive neurons. Scale bar: 5 µm. Calibration bar: grey scale values.**(B)** Quantification of intensity of surface α-SynCAM1 inside dendrites of neurons expressing GFP-R2^WT^, GFP-R2^DEAD^ or GFP-UR2^WT^ normalised to the intensity within dendrites of neurons expressing GFP. Mean intensity values ± SD (a.u.): control (GFP) 1 ± 0.21, GFP-R2^WT^ 0.6 ± 0.18, GFP-R2^DEAD^ 0.86 ± 0.29, GFP-UR2^WT^ 0.88 ± 0.31. N = 4 independent experiments. One-way ANOVA followed by Sidak’s multiple comparisons test. **(C)** Example gSTED image of a neuron expressing HaloTag-UR2-S580 co-stained with postsynaptic marker α-PSD95-S635. HaloTag was labeled using α-HaloTag antibody. Scale bar: 2 µm. Example image and intensity line profile of an individual dendritic spine. Scale bar: 500 nm. **(D)** Example confocal images of surface α-SynCAM1-STAR-RED staining in neurons expressing control shRNA, R2 KD1, or R2 KD2 with re-expression of HaloTag-R2 or HaloTag-UR2. Orange Hot LUT was used to display differences in fluorescence intensity. Dotted circles represent the outlines of PSDs based on corresponding images of α-PSD95-S580. Scale bar: 5 µm. **(E)** Quantification of the intensity of surface α-SynCAM1 inside the PSD of neurons with R2 KD1 and R2 KD1 with re-expression of HaloTag-R2 or HaloTag-UR2, normalised to intensity within the PSD of neurons expressing control shRNA. Mean intensity values ± SD (a.u.): ctrl KD 1.00 ± 0.17, R2 KD1 1.68 ± 0.45, R2 KD1 + R2 0.92 ± 0.36, R2 KD1 + UR2 1.45 ± 0.36. N = 3 independent experiments. One-way ANOVA followed by Sidak’s multiple comparisons test. **(F)** The role of RHBDL2 in regulating synapse architecture and remodelling capacity. The rapid lateral mobility of R2 enables it to scan the neuronal plasma membrane and control abundance of synaptic CAMs that are not engaged in stable adhesion complexes. During synapse remodeling, R2 contributes to dynamic changes in synapse structure by regulating synaptic CAMs that are released from adhesion complexes, thus preventing their reassembly.

We next asked whether the unrestrained mobility of RHBDL2 is required for physiological regulation of synapse composition. Following depletion of endogenous RHBDL2, re-expression of wild-type RHBDL2 restored synaptic SynCAM1 abundance. In contrast, re-expression of immobilised RHBDL2 failed to rescue this phenotype (Figure 6D and 6E), despite its wild type catalytic activity (Figure 5D). Thus, rapid lateral diffusion is essential for RHBDL2 function in neurons, in a manner independent of its proteolytic capacity.

Collectively, these findings demonstrate that the biological activity of RHBDL2 depends not only on its catalytic function, but also on its ability to rapidly scan the neuronal plasma membrane. We propose that RHBDL2 acts as a membrane surveillance protease that continuously explores dendritic and synaptic membranes, selectively removing adhesion molecules that are not incorporated into stable synaptic complexes, while sparing adhesive assemblies engaged within mature synapses. By maintaining the dynamic balance between synaptic stabilisation and adhesion turnover, this lateral diffusion-dependent surveillance mechanism preserves the capacity of mature synapses to undergo structural remodelling and plasticity (Figure 6F).

## Discussion

The ability of synapses to dynamically adjust their structure and molecular composition, and by this their strength, is fundamental to synaptic signalling and plasticity. Our findings identify RHBDL2 as a regulator of synapse organisation and suggest its function extends beyond the control of individual substrates to the broader processing of CAMs at the neuronal surface. Consistent with this, loss of RHBDL2 increases surface SynCAM1 and impairs mGluR-LTD (Figures 2F, 2G, and Figure 3), in agreement with previous observations that increased SynCAM1 expression has similar effects on LTD in a mouse model^39^. Conversely, increased RHBDL2 expression promotes a filopodia-like spine morphology (Figure 1D-1F), which is reminiscent of structural defects observed upon the functional disruption of RHBDL2 substrates ephrin-B2 and ephrin-B3^23,33,34^. Together with our TMD cleavage screen (Figure 2B and 2C), which identified multiple synaptic CAM TMDs as RHBDL2 substrates, these observations support a model in which RHBDL2 regulates the broader synaptic landscape through the collective control of multiple CAMs at the neuronal surface, rather than through the processing of a single dominant substrate.

Our expression data suggest that expression patterns across neuronal development position RHBDL2 to perform such a role (Figure 1A). RHBDL2 expression is very low during early stages of synapse development (DIV7) and increases after synapse formation (DIV19). This temporal pattern correlates with a switch in function of many synaptic CAMs from promoting initial synaptic assembly towards maintaining and dynamically reorganising mature synapses^7–9^. Consistent with a role at established synapses, increasing RHBDL2 expression in mature neurons (DIV16-19) promotes a shift towards more immature, filopodia-like protrusions and alters synapse position (Figure S1E). We therefore speculate that relatively low expression of RHBDL2 during early synaptogenesis favours the establishment of new adhesive contacts, whereas increased RHBDL2 expression in mature neurons provides greater capacity for surveillance and subsequent remodelling of adhesion complexes. RHBDL2 may consequently establish a proteolytic threshold that determines how readily existing synaptic structures can be reorganised, rather than acting as a constitutive destabliser of synapses.

How does RHBDL2 exert such broad effects despite its relatively low abundance? We postulate that its exceptionally rapid mobility at the neuronal surface contributes to its synaptic function (Figure 4). Restricting RHBDL2 mobility impairs its ability to restore correct SynCAM1 distribution without altering its intrinsic proteolytic activity, suggesting that lateral diffusion determines access to functionally relevant substrates (Figures 5 and Figure 6). This is particularly pertinent to neurons, where membrane proteins are segregated across highly specialised compartments^28,50^, and diffusion barriers restrict protein exchange between distinct membrane domains^51,52^. The unusually rapid mobility of RHBDL2 is consistent with a previous study showing that rhomboids can diffuse through membranes more rapidly than proteins with similar sizes and geometries^45^. Studies of bacterial GlpG have linked this behaviour to the unusual architecture of the core rhomboid fold and its interactions with surrounding lipids, which distort the local lipid bilayer and facilitate rapid lateral diffusion^45,53^. Interestingly, RHBDL2 has relatively short cytoplasmic and extracellular regions compared with other rhomboid-family proteins (Figure S4), which likely limit its engagement with cytoskeletal and extra-and intracellular scaffolds that would otherwise restrain lateral mobility. Furthermore, in contrast to other multi-pass synaptic proteins such as GluA1 and mGluR2, it is strictly monomeric^54^, and so its diffusion is unaffected by homo-or heteromeric interactions. We therefore propose that these unique features of RHBDL2 enable its rapid diffusion, which enable it to encounter substrates in all spatially distinct regions of the neuronal plasma membrane, including crowded and selective environments such as the PSD^29,55^.

The combination of broad membrane access and wide substrate repertoire raise a further question: how can RHBDL2 distinguish between CAMs that should be retained within synaptic adhesion complexes from those that should be removed? One possibility is that substrate susceptibility is determined not simply by the identity of the CAM, but by its physical state within the membrane. Synaptic CAMs oligomerise into higher order complexes through interactions between their extracellular domains, TMDs, and intracellular scaffolds^7,8,40,42^. Incorporation into stable adhesion complexes could therefore reduce exposure of a substrate TMD or enhance its stability, whereas unassembled, mislocalised or recently released molecules may become more susceptible to RHBDL2 cleavage. This model is consistent with the established ability of rhomboid proteases to recognise membrane proteins according to the structural properties of their TMDs, rather than through simple sequence motifs^20–22^. As synaptic CAMs exist as multiple isoforms that differ primarily in their extracellular or intracellular domains, this rhomboid-dependent mechanism also provides a means to regulate diverse adhesion molecules within the same family via their common TMDs. The previous observation that bacterial rhomboids cleave membrane subunits of respiratory complexes that fail to become incorporated into functional complexes^56^, indicate that this form of surveillance is evolutionarily widespread.

This model places RHBDL2 within a broader framework of regulators of synaptic weakening and LTD. LTD persists for hours or longer, implying that transient changes in synaptic activity must ultimately be converted into durable alterations in synaptic composition. Current models for LTD largely focus on cytoplasmic post-translational mechanisms involving phosphorylation, ubiquitination, autophagy and endocytic regulation for the regulated removal of glutamate receptors from the postsynapse^2,6,57,58^. These pathways provide important mechanisms for changing receptor abundance, but do not readily explain how trans-synaptic adhesion complexes are remodelled and maintained over the same timescale. We propose that intramembrane proteolysis provides an intrinsically irreversible mechanism for converting a transient change in synaptic state into sustained loss of trans-synaptic adhesive contacts. Such regulation may also address a second, less well-resolved feature of LTD: a need to remodel adhesion proteins within the PSD, which is itself largely devoid of endocytic activity^24^. As described earlier, synaptic CAMs are not distributed uniformly across the neuronal surface, but are incorporated into specialised complexes within the PSD^8^. If release from a stable synaptic complex increases CAM susceptibility to RHBDL2 processing, then cleavage and release of the CAM extracellular domain would prevent its immediate reassociation and favour persistent of the remodelling event. In this model, intramembrane proteolysis by RHBDL2 is not the plasticity signal itself, but it couples the transient induction of LTD to durable reorganisation of synaptic adhesion, complementing known cytoplasmic mechanisms of LTD regulation. This mechanism therefore provides a means of linking the molecular histories of individual synapses to their stability: CAMs that remain appropriately incorporated into functional complexes are retained, whereas those that become disengaged are progressively removed.

Our findings place this RHBDL2-dependent mechanism within the broader landscape of membrane proteolysis in neuronal signalling. Synaptic CAMs are subject to regulation by several other protease families, including ADAM metalloproteases^59–61^, which, like RHBDL2, exhibit broad substrate repertoires and can alter membrane-associated and extracellular signalling^21,35,62^. However, RHBDL2 represents a mechanistically distinct form of proteolytic regulation. ADAM proteases are subject to extracellular and intracellular regulation^63–65^ and, through their soluble extracellular catalytic domains and single-pass membrane topology, are well positioned to access CAMs within oligomeric or higher-order synaptic complexes. ADAM-mediated cleavage may therefore provide a rapid and stimulus-responsive mechanism for the broad remodelling of synaptic protein composition. RHBDL2 activity, in contrast, may provide a more constitutive form of proteostatic control, in which lateral mobility and intramembrane cleavage progressively tune the abundance of surface synaptic CAMs. Evidence for shared substrates for rhomboids and ADAM proteases^21,35,62,66^ raise the possibility that these pathways act in complementary ways, with extracellular shedding and intramembrane proteolysis providing distinct mechanisms for controlling the lifetime and organisation of synaptic membrane proteins.

Altered processing of substrates by other intramembrane proteases γ-secretase and SPPL2b are implicated in neuropathology^14,16^. RHBDL2 has not been studied in this context, as its neuronal function was unknown. That RHBDL2 regulates synapse stability raises the possibility that dysregulation of rhomboids may also contribute to neurological disease. For instance, any reduction in RHBDL2 levels or lateral mobility could shift mature synapses towards excessive stability, which could lead to substantial defects in neuronal signalling. Additionally, genetic disruption of the related and neuronally enriched orphan rhomboids RHBDL1 and RHBDL3 are reported to display embryonic lethality^67^, which places them as potential regulators of neuronal signaling. Therefore, determining how other intramembrane proteases coordinate lateral membrane organisation with neuronal function will likely provide important insights into neurological disorders.

More generally, polarised cell types such as neurons, epithelia and immune cells share a common requirement for stable intercellular signalling complexes, such as those at synapses and cell-cell junctions. These complexes must all retain the capacity to remodel in response to changing demands^68,69^. Therefore diffusion-dependent proteolytic regulation should be explored as a widespread mechanism that shapes the signalling and architecture of cell-cell interfaces.

## Materials and methods

### Animals

All animal care and experimental procedures were conducted in accordance with the regulations and guidelines of University of Bristol and the UK Home Office.

### Antibodies and reagents

The following primary antibodies were used in this study: mouse anti-AMPAR (Synaptic System, #182411, RRID AB_2619876, 1:100 dilution), mouse anti-Bassoon (Enzo, #ADI-VAM-PS003-F, RRID AB_10618753, 1:1000 dilution), rabbit anti-GFP (MBL, #598, RRID AB_591819, 1:1000 dilution), rabbit anti-HaloTag (Promega, #G9281, RRID AB_713650, 1:500 dilution), rabbit anti-Homer1 (Synaptic System, #160003, RRID AB_887730, 1:500 dilution), mouse anti-PSD95 (Neuromab, #75–028, RRID AB_2307331, 1:300 dilution), nanobody FluoTag-X2 anti-PSD95-STAR635P (NanoTag Biotechnologies, #N3702-AB635P-L, RRID AB_3076102, 1:200 dilution), chicken anti-SynCAM1 (MGL, #CM004-3, RRID AB_592783, 1:750 dilution). The following secondary antibodies were used (all 1:200 dilution): goat Alexa Fluor 488-conjugated anti-rabbit IgG (Invitrogen, #A-11034), goat STAR580-conjugated anti-rabbit IgG (Abberior, #ST580-1002-500UG) and anti-mouse IgG (Abberior, # ST580-1001-500UG), goat STAR635P-conjugated anti-rabbit IgG (Abberior, #ST635P-1002-500UG) and anti-mouse (Abberior, #ST635P-1001-500UG), goat Abberior STAR RED-conjugated anti-chicken IgY (Abberior, #STRED-1005-500UG). The HaloTag ligands JFX650 and JF635i used in this study were a gift from the Lavis lab and Open Chemistry team (Janelia). The following chemical reagents were used: (S)-3,5-DHPG (HelloBio, #HB0045), Latrunculin B (Merck, #428020), PNPP substrates tablets (Thermo Scientific, #34047), Pierce diethanolamine substrate buffer 5x (Thermo Scientific, #34064), Tetrodotoxin citrate (Torcris Bioscience, #1069).

### DNA plasmids

The following plasmids were described previously: pAAV-hSyn-GCaMP6f (Addgene plasmid #100837,^30^); pEGFP-AP-L1-CAM-TMD, pEGFP-AP-NCAM-TMD, pEGFP-AP-NCad-TMD, pEGFP-AP-Nlg3-TMD, pEGFP-AP-Calnexin-TMD ^21^; pcDNA3.1 (Invitrogen); psPAX2, pMD2.G ^70^. Plasmid pXLG-shRNA-FF3 containing control shRNA against firefly luciferase (shRNA sequence 5’CCGCTGAATTGGAATCCTT3’) was a gift from Kevin Wilkinson, University of Bristol. pLVX-hSyn vector was generated by exchanging the CMV promotor in pLVX (Takara) to the human synapsin promotor (hSyn) from pAAV-hSyn-GCaMP6f. The DNA sequence of rat RHBDL1, RHBDL2 and RHBDL3 was obtained as a gBlock (Integrated DNA Technologies) and used as PCR template for further cloning into pcDNA3.1 (for rhomboid expression in cell lines) and pLVX-hSyn (for expression in neurons) vectors. The sequence of R2^DEAD^ was created by site-directed mutagenesis (QuikChange Mutagenesis kit, Agilent) by mutating serine at position 185 to alanine. pEGFP (Clontech) was used as a PCR template to generate all plasmids containing GFP. TUBB5-HaloTag (Addgene plasmid # 64691,^71^) was used as PCR template to generate all plasmids containing HaloTag. Plasmids pcDNA-HaloTag-R1, pcDNA-HaloTag-R2, pcDNA-HaloTag-R2^DEAD^, pcDNA-HaloTag-R3, pLVX-hSyn-GFP, pLVX-hSyn-GFP-R2, pLVX-hSyn-HaloTag-R2, pLVX-hSyn-GFP-R2^DEAD^, pLVX-hSyn-GFP-R3 were cloned using In-Fusion cloning (In-Fusion Snap Assembly, Takara). cDNA coding for SPPL2b, a gift from Colin Adrain, Queen’s University Belfast, was used as a PCR template to generate pcDNA-HaloTag-SPPL2b. pLEX-iRhom1-3xHA ^72^ was use as PCR template for cloning pcDNA-HaloTag-iRhom1. Plasmid pRK5-HaloTag-mGluR2 was generated by replacing SEP with HaloTag in pRK5-SEP-mGluR2 ^73^. The plasmid YFP-GT46 ^74^ was used as PCR template to generate pcDNA-HaloTag-GT46 and pLVX-hSyn-HaloTag-GT46. pEGFP-GluA1 ^75^ was used as PCR template to create pLVX-hSyn-HaloTag-GluA1. Knock-down vectors pXAG-shRNA-R2-KD1 (with shRNA sequence 5’GGATTGTCTCCAAGTGGAT3’), pXLG-shRNA-R2-KD2 (with shRNA sequence 5’GCCTGGAGGTTTATCTCAT3’) and pXLG-shRNA-R3-KD (with shRNA sequence 5’TTGACCGCAAGTGGTACTATG 3’) were cloned according to a previously published protocol ^70^. Plasmids pXAG-shRNA-R2-KD1-hSyn-HaloTag-R2, pXLG-shRNA-R2-KD2-hSyn-GFP-R2, pXLG-shRNA-FF3-hSyn-GCamp6F and pXAG-shRNA-R2-KD1-hSyn-GCaMP6f were generated by replacing SFFV promotors and eGFP with hSyn-HaloTag-R2 from pLVX-hSyn-HaloTag-R2, hSyn-GFP-R2 from pLVX-hSyn-GFP-R2 and hSyn-GCaMP6f from pAAV-hSyn-GCaMP6f, respectively. pGEM-SNAPf-UtropinCH ^76^, a gift from Binyam Mogessie, Yale School of Medicine, was used as PCR template for the calponin-homology domain of utrophin to clone the vectors pcDNA-HaloTag-UR2, pcDNA-HaloTag-UR2^DEAD^, pLVX-hSyn-GFP-UR2, pLVX-HaloTag-UR2, pLVX-GFP-UtrophinCH, and pXLG-shRNA-R2-KD1-hSyn-HaloTag-UR2. pEGFP-AP-SynCAM1_TMD was cloned as described previously ^21^ using the following oligonucleotides for SynCAM1_TMD FOR:5’GGTGCTGGTGCTGTCGACATTGGGGCCGTCGATCACGCCGTGATCGGCGGCGTTGTTGCCGTGGTTGTCTTTGCTATGCT3’ and REV: 5’GATTATGATCTAGAGTCGCGGCCGCTCACCTGGCAAAATATCTGCCCAGGATAATAAGGAGGCACAGCATAGCAAAGACAACCA3’. Plasmids pCMV6-Efnb2 and pCMV6-Efnb3 (OriGene Technologies) were used as a PCR template for the TMD region of ephrin-B2 in pEGFP-AP-ephrin-B2_TMD and ephrin-B3 in pEGFP-AP-ephrin-B3_TMD. All sequences were verified by Sanger sequencing (Source Genomics, Cambridge).

### Primary rat neuronal culture and transfection

Dissociated hippocampal and cortical cultures from embryonic day 18 (E18) Wister rat brains (Charles River Laboratories) of both genders were performed as described previously^77^. Hippocampal neurons were plated on round 18-mm glass coverslips (thickness 1.5) coated with poly-D-lysine (for SMT experiments) or poly-L-lysine (for all other experiments) (1 mg/ml in borate buffer, Sigma-Aldrich) at a density of 7.5×10^4^ (for SMT experiments) or 1×10^5^ cells (for all other experiments) per well in 12-well plate. Cortical neurons were plated in 6-well plates (for KD experiments) or in 35 mm dishes (for mRNA time course experiment) coated with poly-L-lysine (0.5 mg/ml in borate buffer) at density 5×10^5^ cells per well or dish. After dissociation, neurons were plated in plating medium (Neurobasal Medium (Gibco) supplemented with 2% B27 (Gibco), 10% horse serum (Sigma-Aldrich), 2 mM GlutaMAX (Gibco) and 1% penicillin/streptomycin (Gibco)). 2h after plating, plating medium was replaced with feeding medium (Neurobasal Medium supplemented with 2% B27, 1 mM GlutaMAX and 0.5% penicillin/streptomycin). Starting from DIV7, once per week, half the conditioned medium was exchanged for fresh feeding medium, or feeding-plus medium (Neurobasal Plus Medium (Gibco) supplemented with 2% B27 Plus (Gibco), 1 mM GlutaMAX; for all experiments that required surface IF). Neurons were transfected at DIV9 or DIV16 using Lipofectamine 2000 (Invitrogen). All DNA used for neuronal transfection was purified using plasmid midi-prep kit (Qiagen). Shortly before transfection, conditioned medium was collected and replaced with fresh feeding medium. After 30 min incubation at RT, a mixture of 2 µg of DNA with 3.3. µl Lipofectamine per well of 12-well plate was added to neurons. Neurons were incubated with DNA/Lipofectamine for 1h at 37°C, 5% CO_2_. Next, neurons were briefly washed with fresh Neurobasal medium and conditioned medium was placed back onto wells. All subsequent experiments were performed at DIV19-20.

### HEK293T culture and U2OS culture and transfection

HEK293T (ATCC CRL-3216) and U2OS cells (ATCC HTB-96) were cultured in high-glucose DMEM (Sigma-Aldrich) supplemented with 10% fetal calf serum (Sigma-Aldrich), 1% glutamine (Gibco) and 1% penicillin/streptomycin (Gibco). 24h before transfection U2OS cells were seeded on round 18-mm glass coverslips (thickness 1.5) in 12-well plates. U2OS cells were transfected 48h before experiments, using FuGene6 HD transfection reagent (Promega). A mixture of 200 ng DNA with 3.3 µl FuGene6 in 200 µl Opti-MEM medium (Gibco) per well was incubated at RT for 30 min and subsequently added to cells.

### Lentivirus production and transduction

Lentivirus production was performed using HEK293T cells, which were seeded at a density of 7×10^6^ cells per 10 cm dish 24h before transfection. Per 10 cm dish, a total 20 µg of DNA of packagaing plasmids psPAX2 and pMD2.G with the shRNA containing pXLG vector (in 1:1:1 a stoichometric ratio) were mixed with 40 µl of PEI (Polysciences) in 500 µl Opti-MEM medium. After 30 min incubation at RT, the mixture of DNA and PEI was added to cells. 6h after transfection, medium was exchanged and for the following 60h lentivirus was secreted into 7 ml of collection medium (3.5 ml of supplemented DMEM and 3.5 ml of Opti-MEM). Next, supernatants was collected, briefly centrifuged and then filtered using 0.45 µm pore size filter to remove cell debris. Cleared supernatant was concentrated using Amicon Ultra-4 100K columns (Millipore) to ∼50 µl. Fresh, concentrated supernatant containing lentivirus particles was added at DIV9 to hippocampal neurons (7-15 µl per well in 12-well plate) or cortical neurons (30-45 µl per well in 6-well plate). Due to the low expression levels of RHBDL2 in knock-down/rescue experiments, HaloTag-RHBDL2 and GFP-RHBDL2 were labeled with α-HaloTag or α-GFP antibody respectively, and secondary antibody Alexa Fluor 488. Lentivirus infection was used for all knock-down experiments except the confocal imaging in Figure S1C and S1D, and GCaMP imaging in Figure S2B.

### AP-TMD shedding assay

The AP-TMD shedding assay was performed using HEK293T cells as described previously^21^ with some modifications. In brief, 24h before transfection, cells were seeded at a density of 1.5×10^5^ per well of 24-well plate coated with poly-L-lysine (0.1 mg/ml, R&D Systems). HEK293T cells were co-transfected with plasmid containing AP-substrate-TMD and pcDNA3.1 vector (control) or pcDNA3.1 vector containing HaloTag-tagged R2^WT^ or R2^DEAD^ (150 ng DNA per plasmid per well) using FuGene6 HD transfection reagent (1.5 µl per well; Promega). 24h after transfection, culture medium was exchanged for 300 µl of Opti-MEM without phenol red (Gibco). 16h later, detection of AP activity was performed. To assess levels of AP activity, culture supernatant was collected and cells were lysed with lysis buffer (1% Triton X-100, 150 mM NaCl, 50 mM Tris-HCl, pH7.4, supplemented with protease inhibitor cocktail (Roche)). Both supernatant and cell lysates were cleared by brief centrifugation. Next, undiluted supernatants and cell lysates diluted 1:5 in water were added to equal volumes of PNPP substrate, incubated at RT for 10 min and absorbance at 405 nm was measured using a plate reader. Analysis of AP-TMD shedding results was performed as described previously^21^.

### qRT-PCR

RNA was isolated from cortical neurons using RNeasy Mini kit (Qiagen) following the manufacturer’s instructions. 1.1 µg of RNA per condition was used for synthesis of cDNA using Super Script III First-Strand Synthesis Super Mix (Invitrogen) following the manufacturer’s instruction. Next, cDNA was diluted 1:2 in water and qRT-PCR was performed using TaqMan gene expression assay (Applied Bioscience) and QuantStudio 3 system (Applied Bioscience). The following TaqMan probes were used for detection of rhomboids: *RHBDL2* Rn01515733-m1, *RHBDL3* Rn01750781-m1, and for detection of housekeeping genes: *GAPDH* Rn01775763-g1 and *Actb* Rn00667869_m1. Relative mRNA levels were calculated based on ΔΔCt values.

### Immunostaining, confocal and gSTED imaging

Neurons at DIV19 were fixed with pre-warmed 4% PFA (Thermo Scientific) with 4% sucrose (Fisher Chemicals) for 15 min at RT. Next, cells were washed three times for 10 min with PBS supplemented with 100 mM glycine (Severn Biotech). Neurons were permeabilised and blocked with 0.1% Triton-×100 (Sigma-Aldrich), 10% natural goat serum (Invitrogen) and 100 mM glycine in PBS for 1h at 37°C. Neurons were incubated with primary antibodies diluted in PBS supplemented with 5% natural goat serum, 0.1% Triton X-100 and 100 mM glycine overnight at 4^0^C. Next, cells were washed three times for 10 min with PBS supplemented with 100 mM glycine, and then incubated with secondary antibodies diluted in PBS supplemented with 5% natural goat serum, 0.1% Triton X-100 and 100 mM glycine 2h at RT (for confocal) or overnight at 4^0^C (for gSTED). Neurons were washed twice for 10 min with PBS with 100 mM glycine, followed by two 10 min washes with PBS and mounted in ProLong Diamond Antifade mounting medium (Invitrogen) on glass slides. Confocal imaging was performed with Leica SP8 equipped with a HC PL APO CS2 63/1.4 oil objective, and controlled by LASX software. GFP and Alexa Fluor 488 were excited with a 488 nm laser line of Ar laser, Abberior STAR580 with a 561 nm DPSS laser, and Abberior STAR635P and STAR RED with a 633 nm Red He/Ne laser. Fluorescence emission was detected using Leica HyD detectors. Dual-color gated STED imaging was performed with Leica SP8X STED 3X, using a HCX PL APO 100/1.4 oil STED objective. Abberior STAR580 was excited using a 561 nm laser line, and Abberior STAR635P and STAR RED were excited using a 633 nm laser line (pulsed white light laser, 80 MHz). All dyes used for STED imaging were depleted using a 775 nm pulsed depletion laser. Fluorescence emission was detected using Leica HyD detectors with gating from 0.6 ns to 6 ns and a pinhole size of 0.85 AU.

### Chemical LTD protocol and surface immunostaining

To induce chemical LTD, 50 µM (S)-3,5-DHPG was added to conditioned medium of neurons at DIV19 for 30 min. Next, surface IF was performed. For all experiments requiring surface labeling, the following protocol was used. Live neurons were incubated with primary antibody against extracellular epitopes of AMPAR or SynCAM1, diluted in modified Tyrode’s buffer (25 mM HEPES, 119 mM NaCl, 2.4 mM KCl, 2 mM CaCl_2_, 2 mM MgCl_2_, 30 mM glucose, pH 7.4) and supplemented with 10% of BSA for 10 min on ice. Next, cells were washed once with ice-cold Tyrode’s buffer and fixed with 4% PFA with 4% sucrose for 15 min. Neurons underwent three 10 min washes with PBS with 100 mM glycine, and were then blocked with 10% natural goat serum and 100 mM glycine in PBS for 1h at 37°C. Neurons were incubated with secondary antibody diluted in PBS supplemented with 5% natural goat serum and 100 mM glycine for 4h at RT. Neurons were washed three times with PBS with 100 mM glycine for 10 min. Next, cells were permeabilised and blocked with blocking solution containing 0.1% Triton X-100 for 1h at 37°C. All subsequent steps followed the protocol of immunostaining described above.

### Single molecule tracking (SMT) imaging

20 min before the start of SMT experiments, a picomolar concentration of HaloTag ligand was added to the conditioned medium of neurons or U2OS cells. Next, cells were washed and mounted in an imaging chamber (Warner Instruments). SMT was carried out in modified Tyrode’s buffer with an Abbelight SAFe 350 system based on an Olympus IX83 microscope equipped with a 100x/1.5 oil TIRF objective and controlled by Abbelight NEO software. 638 nm diode lasers were used to excite the HaloTag ligands. Fluorescence emission was detected using a sCMOS camera (Hamamatsu ORCA-Fusion). 2000 frames for SMT in neurons, and 3000 frames for SMT in U2OS, were acquired in stream mode at 40Hz in TIRF. All imaging was performed at 37°C (using a PECON environmental chamber). In SMT experiments with utrophin-RHBDL2 (Figure 5C), neurons were incubated 2 – 5 min with 1 µM Latrunculin B (LatB).

### Ca^2+^ imaging using GCaMP6f

GCaMP imaging was carried out in modified Tyrode’s buffer without Mg^2+^ (25 mM HEPES, 121 mM NaCl, 2.4 mM KCl, 2 mM CaCl2, 30 mM glucose, pH 7.4), supplemented with 3 µM TTX with an Abbelight SAFe 350 system based on an Olympus IX83 microscope equipped with a 100x/1.5 oil TIRF objective and controlled by Abbelight NEO software. A blue LED diode was used to excite GCaMP, and a 638 nm diode laser was used to excite the HaloTag ligand JFX650 using epifluorescence illumination. Fluorescence emission was detected using sCMOS cameras (Hamamatsu ORCA-Fusion). 3000 frames (for GCaMP channel) or 30 frames (for HaloTag channel) in stream mode at 20Hz were acquired. All imaging was performed at 37°C (using a PECON environmental chamber).

## Data Analysis

### Quantification of surface intensity of AMPAR and SynCAM1

For quantification of the intensity of surface AMPAR or SynCAM1 inside the PSD in KD experiments, PSD masks were created based on α-PSD95 images using the Spot Detector plug-in in the Icy bioimaging platform^78^. PSDs labeled with the antibody α-PSD95 (for SynCAM1 analysis) were detected as spots with a size of ∼7 pixels with sensitivity 100, and ∼13 pixels with sensitivity 110. PSDs labeled with the nanobody α-PSD95 (for AMPAR analysis) were detected as spots with size ∼7 pixels with sensitivity 85, and ∼13 pixels with sensitivity 100. Based on these detections, binary images were created. Next, binary images were loaded into ImageJ^79^ and spots in the binary mask were detected and saved as ROIs. ROIs were applied to images of AMPAR or SynCAM1 and mean intensity values were calculated for each ROI. For quantification of the intensity of total surface SynCAM1 in OE experiments, a GFP mask was created using ImageJ. The GFP image was blurred with Gaussian blur (radius 3) and an auto intensity threshold was applied to create an outline of the GFP expressing cell. Next, this outline was applied to SynCAM1 images and mean intensity values were calculated. For quantification of the intensity of surface SynCAM1 inside the PSD in OE experiments, first a GFP mask was created to provide an outline of a transfected cell. Next, this outline was applied to the PSD mask to create two new masks with PSD spots inside transfected neurons, and also those outside transfected neurons, which were used for further analysis. All images used in these analyses were the maximum intensity projections of the Z-stacks.

### GCaMP Ca^2+^ transient analysis

Detection of Ca^2+^ transients was performed as described previously^80^ with some minor modifications using a workflow implemented into MIA v1.6.0 ImageJ plug-in^81^ (https://mianalysis.github.io/). In short, a stack of images was loaded into the MIA workflow and ROIs were manually drawn around all in focus spines that were visibly separated from shaft, and line ROIs were drawn along in focus dendritic shafts. Next, a median rolling threshold was applied and all events inside ROIs with an intensity that exceeded two standard deviations of the mean signal was calculated, based on 40 frames prior to event detection. For each detected event, an F/F_0_ trace was calculated. For the final frequency analysis, events were filtered based on minimum initial gradient (> 40) and minimum peak amplitude (> 0.035). Frequency of Ca^2+^ transients in spines (per minute) was calculated by dividing the number of detected events by the number of all selected spine ROIs, regardless of their activity. Frequency of Ca^2+^ transients in dendritic shaft (per minute) was calculated by dividing the number of all detected shaft events by the total length of linear ROIs and normalised to 50 µm.

### Single molecule tracking (SMT) analysis

All analysis of SMT experiments was performed as described previously^73^ with some modifications. Detection and Gaussian fitting of molecules were performed using the DoM v1.2.5 ImageJ plug-in (https://github.com/UU-cellbiology/DoM_Utrecht), with an intensity threshold of 3.0 SNR and detection of PSD SD 1.7 pixel. Only molecules with localisation precision of < 50 nm were used for analysis. Tracking, calculation of diffusion coefficient (D_inst_) and mobile fraction were performed using custom-written Matlab (MathWork) scripts described previously^46,82^ (https://github.com/ManonWestra/SMT_analysis). Only trajectories longer than 25 frames were used to calculate the instantaneous diffusion coefficient, which was estimated by fitting the slope through the four initial points of the MSD curves. Only fields of view with minimum 100 reconstructed trajectories were used for further analysis. Classification of mobile/immobile trajectories was based on the ratio calculated using formula 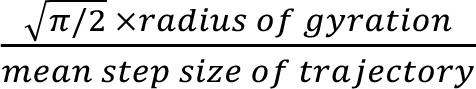. All trajectories with a ratio of < 2.11 were classified as immobile.

### Statistical analysis

All statistical analysis was performed using GraphPad Prism 10. All statistical tests used in this study and the number of experimental replicates are described in the figure legends. In all figures, *P* value < 0.05 was indicated with *, P < 0.01 with ** and P <0.001 with ***. All graphs were created using GraphPad Prism 10. All figures were generated in Inkscape.

## Acknowledgements

We thank Harold MacGillavry, Vrije Universiteit Amsterdam, and Jack Mellor, University of Bristol, for critical reading of the manuscript. We thank Yasuko Nakamura, University of Bristol, for a helpful discussion. We thank Stephen Cross from the Wolfson Bioimaging Facility, University of Bristol, for writing the MIA workflow to analyse GCaMP data and Dominic Alibhai from the Wolfson Bioimaging Facility, University of Bristol for microscopy support. We acknowledge the Wolfson Bioimaging Facility, University of Bristol, for access and support with the Abbelight SAFe 360 funded by a BBSRC Alert 19 equipment grand (BB/T017597/1) and with the Leica SP8. We thank Jana Koth and Cyril Lai from the Wolfson Imaging Centre, MRC WIMM, University of Oxford, for access and support with the Leica TCS SP8 STED 3X. This research was supported by following grants: a BBSRC Research Grant (BB/Y009649) to AB and AGG, and a Sir Henry Dale fellowship jointly funded by the Wellcome Trust and Royal Society (221784/Z/20/Z) to AGG.

## Author Contributions

Conceptualisation: AB, AGG; Investigation: AB, AGG; Methodology: AB, AGG; Supervision: AGG; Visualisation: AB; Writing: AB, AGG.

## References

1. Choquet, D., and Triller, A. (2003). The role of receptor diffusion in the organization of the postsynaptic membrane. Nat. Rev. Neurosci. 4, 251–265. 10.1038/nrn1077.

2. Citri, A., and Malenka, R.C. (2008). Synaptic Plasticity: Multiple Forms, Functions, and Mechanisms. Neuropsychopharmacology 33, 18–41. 10.1038/sj.npp.1301559.

3. Compans, B., Camus, C., Kallergi, E., Sposini, S., Martineau, M., Butler, C., Kechkar, A., Klaassen, R. V., Retailleau, N., Sejnowski, T.J., et al. (2021). NMDAR-dependent long-term depression is associated with increased short term plasticity through autophagy mediated loss of PSD-95. Nat. Commun. 12, 2849. 10.1038/s41467-021-23133-9.

4. Kallergi, E., Daskalaki, A.-D., Kolaxi, A., Camus, C., Ioannou, E., Mercaldo, V., Haberkant, P., Stein, F., Sidiropoulou, K., Dalezios, Y., et al. (2022). Dendritic autophagy degrades postsynaptic proteins and is required for long-term synaptic depression in mice. Nat. Commun. 13, 680. 10.1038/s41467-022-28301-z.

5. Huganir, R.L., and Nicoll, R.A. (2013). AMPARs and Synaptic Plasticity: The Last 25 Years. Neuron 80, 704–717. 10.1016/j.neuron.2013.10.025.

6. Diering, G.H., and Huganir, R.L. (2018). The AMPA Receptor Code of Synaptic Plasticity. Neuron 100, 314–329. 10.1016/j.neuron.2018.10.018.

7. Dalva, M.B., McClelland, A.C., and Kayser, M.S. (2007). Cell adhesion molecules: Signalling functions at the synapse. Nat. Rev. Neurosci. 8, 206–220. 10.1038/nrn2075.

8. Missler, M., Südhof, T.C., and Biederer, T. (2012). Synaptic cell adhesion. Cold Spring Harb. Perspect. Biol. 4, 1–18. 10.1101/cshperspect.a005694.

9. Verpoort, B., and de Wit, J. (2024). Cell Adhesion Molecule Signaling at the Synapse: Beyond the Scaffold. Cold Spring Harb. Perspect. Biol. 16, a041501. 10.1101/cshperspect.a041501.

10. Südhof, T.C. (2018). Towards an Understanding of Synapse Formation. Neuron 100, 276–293. 10.1016/j.neuron.2018.09.040.

11. Biederer, T., Kaeser, P.S., and Blanpied, T.A. (2017). Transcellular Nanoalignment of Synaptic Function. Neuron 96, 680–696. 10.1016/j.neuron.2017.10.006.

12. Strisovsky, K. (2013). Structural and mechanistic principles of intramembrane proteolysis – lessons from rhomboids. FEBS J. 280, 1579–1603. 10.1111/febs.12199.

13. Kühnle, N., Dederer, V., and Lemberg, M.K. (2019). Intramembrane proteolysis at a glance: From signalling to protein degradation. J. Cell Sci. 132. 10.1242/JCS.217745.

14. Badman, J., Parracino, A., Kumar, R., and Tambaro, S. (2025). Insights into the intramembrane protease SPPL2b and its substrates: Functions and disease implications. Sci. Signal. 18, 1–14. 10.1126/scisignal.adt2272.

15. Malvankar, S.R., and Wolfe, M.S. (2026). The γ-secretase complex: from discovery to a therapeutic target. RSC Chem. Biol. 7, 1456–1481. 10.1039/D6CB00022C.

16. De Strooper, B., and Chávez Gutiérrez, L. (2015). Learning by Failing: Ideas and Concepts to Tackle γ-Secretases in Alzheimer’s Disease and Beyond. Annu. Rev. Pharmacol. Toxicol. 55, 419–437. 10.1146/annurev-pharmtox-010814-124309.

17. Annaert, W., and De Strooper, B. (2002). A Cell Biological Perspective on Alzheimer’s Disease. Annu. Rev. Cell Dev. Biol. 18, 25–51. 10.1146/annurev.cellbio.18.020402.142302.

18. Freeman, M. (2014). The Rhomboid-Like Superfamily: Molecular Mechanisms and Biological Roles. Annu. Rev. Cell Dev. Biol. 30, 235–254. 10.1146/annurev-cellbio-100913-012944.

19. Freeman, M. (2008). Rhomboid Proteases and their Biological Functions. Annu. Rev. Genet. 42, 191–210. 10.1146/annurev.genet.42.110807.091628.

20. Moin, S.M., and Urban, S. (2012). Membrane immersion allows rhomboid proteases to achieve specificity by reading transmembrane segment dynamics. Elife 2012, 1–16. 10.7554/eLife.00173.

21. Grieve, A.G., Yeh, Y.C., Chang, Y.F., Huang, H.Y., Zarcone, L., Breuning, J., Johnson, N., Stříšovský, K., Brown, M.H., Parekh, A.B., et al. (2021). Conformational surveillance of Orai1 by a rhomboid intramembrane protease prevents inappropriate CRAC channel activation. Mol. Cell 81, 4784–4798.e7. 10.1016/j.molcel.2021.10.025.

22. Urban, S., and Freeman, M. (2003). Substrate Specificity of Rhomboid Intramembrane Proteases Is Governed by Helix-Breaking Residues in the Substrate Transmembrane Domain. Mol. Cell 11, 1425–1434. 10.1016/S1097-2765(03)00181-3.

23. Pascall, J.C., and Brown, K.D. (2004). Intramembrane cleavage of ephrinB3 by the human rhomboid family protease, RHBDL2. Biochem. Biophys. Res. Commun. 317, 244–252. 10.1016/j.bbrc.2004.03.039.

24. Blanpied, T.A., Scott, D.B., and Ehlers, M.D. (2002). Dynamics and Regulation of Clathrin Coats at Specialized Endocytic Zones of Dendrites and Spines. Neuron 36, 435–449. 10.1016/S0896-6273(02)00979-0.

25. Catsburg, L.A.E., Westra, M., van Schaik, A.M.L., and Macgillavry, H.D. (2022). Dynamics and nanoscale organization of the postsynaptic endocytic zone at excitatory synapses. Elife 11, 1–23. 10.7554/eLife.74387.

26. Li, T.P., and Blanpied, T.A. (2016). Control of transmembrane protein diffusion within the postsynaptic density assessed by simultaneous single-molecule tracking and localization microscopy. Front. Synaptic Neurosci. 8, 1–14. 10.3389/fnsyn.2016.00019.

27. MacGillavry, H.D., Song, Y., Raghavachari, S., and Blanpied, T.A. (2013). Nanoscale scaffolding domains within the postsynaptic density concentrate synaptic ampa receptors. Neuron 78, 615–622. 10.1016/j.neuron.2013.03.009.

28. MacGillavry, H.D., Kerr, J.M., and Blanpied, T.A. (2011). Lateral organization of the postsynaptic density. Mol. Cell. Neurosci. 48, 321–331. 10.1016/j.mcn.2011.09.001.

29. Renner, M.L., Cognet, L., Lounis, B., Triller, A., and Choquet, D. (2009). The excitatory postsynaptic density is a size exclusion diffusion environment. Neuropharmacology 56, 30–36. 10.1016/j.neuropharm.2008.07.022.

30. Chen, T.-W., Wardill, T.J., Sun, Y., Pulver, S.R., Renninger, S.L., Baohan, A., Schreiter, E.R., Kerr, R.A., Orger, M.B., Jayaraman, V., et al. (2013). Ultrasensitive fluorescent proteins for imaging neuronal activity. Nature 499, 295–300. 10.1038/nature12354.

31. Grabrucker, A., Vaida, B., Bockmann, J., and Boeckers, T.M. (2009). Synaptogenesis of hippocampal neurons in primary cell culture. Cell Tissue Res. 338, 333–341. 10.1007/s00441-009-0881-z.

32. Körber, N., and Stein, V. (2016). In vivo imaging demonstrates dendritic spine stabilization by SynCAM 1. Sci. Rep. 6, 24241. 10.1038/srep24241.

33. Bissen, D., Kracht, M.K., Foss, F., Hofmann, J., and Acker-Palmer, A. (2021). EphrinB2 and GRIP1 stabilize mushroom spines during denervation-induced homeostatic plasticity. Cell Rep. 34, 108923. 10.1016/j.celrep.2021.108923.

34. Segura, I., Essmann, C.L., Weinges, S., and Acker-Palmer, A. (2007). Grb4 and GIT1 transduce ephrinB reverse signals modulating spine morphogenesis and synapse formation. Nat. Neurosci. 10, 301–310. 10.1038/nn1858.

35. Johnson, N., Březinová, J., Stephens, E., Burbridge, E., Freeman, M., Adrain, C., and Strisovsky, K. (2017). Quantitative proteomics screen identifies a substrate repertoire of rhomboid protease RHBDL2 in human cells and implicates it in epithelial homeostasis. Sci. Rep. 7, 7283. 10.1038/s41598-017-07556-3.

36. Battistini, C., Rehman, M., Avolio, M., Arduin, A., Valdembri, D., Serini, G., and Tamagnone, L. (2019). Rhomboid-like-2 intramembrane protease mediates metalloprotease-independent regulation of cadherins. Int. J. Mol. Sci. 20. 10.3390/ijms20235958.

37. Baker, R.P., Wijetilaka, R., and Urban, S. (2006). Two Plasmodium Rhomboid Proteases Preferentially Cleave Different Adhesins Implicated in All Invasive Stages of Malaria. PLoS Pathog. 2, e113. 10.1371/journal.ppat.0020113.

38. Fogel, A.I., Stagi, M., Perez De Arce, K., and Biederer, T. (2011). Lateral assembly of the immunoglobulin protein SynCAM 1 controls its adhesive function and instructs synapse formation. EMBO J. 30, 4728–4738. 10.1038/emboj.2011.336.

39. Robbins, E.M., Krupp, A.J., Perez de Arce, K., Ghosh, A.K., Fogel, A.I., Boucard, A., Südhof, T.C., Stein, V., and Biederer, T. (2010). SynCAM 1 Adhesion Dynamically Regulates Synapse Number and Impacts Plasticity and Learning. Neuron 68, 894–906. 10.1016/j.neuron.2010.11.003.

40. Fogel, A.I., Akins, M.R., Krupp, A.J., Stagi, M., Stein, V., and Biederer, T. (2007). SynCAMs Organize Synapses through Heterophilic Adhesion. J. Neurosci. 27, 12516–12530. 10.1523/JNEUROSCI.2739-07.2007.

41. Perez de Arce, K., Schrod, N., Metzbower, S.W.R., Allgeyer, E., Kong, G.K.-W.K.W., Tang, A.-H.H., Krupp, A.J., Stein, V., Liu, X., Bewersdorf, J., et al. (2015). Topographic Mapping of the Synaptic Cleft into Adhesive Nanodomains. Neuron 88, 1165–1172. 10.1016/j.neuron.2015.11.011.

42. Benson, D.L., Schnapp, L.M., Shapiro, L., and Huntley, G.W. (2000). Making memories stick: Cell-adhesion molecules in synaptic plasticity. Trends Cell Biol. 10, 473–482. 10.1016/S0962-8924(00)01838-9.

43. Palmer, M.J., Irving, A.J., Seabrook, G.R., Jane, D.E., and Collingridge, G.L. (1997). The group I mGlu receptor agonist DHPG induces a novel form of LTD in the CA1 region of the hippocampus. Neuropharmacology 36, 1517–1532. 10.1016/S0028-3908(97)00181-0.

44. Clifton, B.R., Corey, R.A., and Grieve, A.G. (2025). Structural and energetic insights into human rhomboid proteases reveal a unique lateral gating mechanism for orphan family members, 10.1101/2025.11.21.689725 https://doi.org/10.1101/2025.11.21.689725.

45. Kreutzberger, A.J.B., Ji, M., Aaron, J., Mihaljević, L., and Urban, S. (2019). Rhomboid distorts lipids to break the viscosity-imposed speed limit of membrane diffusion. Science (80-.). 363, 1–23. 10.1126/science.aao0076.

46. Westra, M., and MacGillavry, H.D. (2022). Precise Detection and Visualization of Nanoscale Temporal Confinement in Single-Molecule Tracking Analysis. Membranes (Basel). 12, 650. 10.3390/membranes12070650.

47. Winder, S.J., Hemmings, L., Maciver, S.K., Bolton, S.J., Tinsley, J.M., Davies, K.E., Critchley, D.R., and Kendrick-Jones, J. (1995). Utrophin actin binding domain: analysis of actin binding and cellular targeting. J. Cell Sci. 108, 63–71. 10.1242/jcs.108.1.63.

48. Melak, M., Plessner, M., and Grosse, R. (2017). Actin visualization at a glance. J. Cell Sci. 130, 525–530. 10.1242/jcs.189068.

49. Hotulainen, P., and Hoogenraad, C.C. (2010). Actin in dendritic spines: connecting dynamics to function. J. Cell Biol. 189, 619–629. 10.1083/jcb.201003008.

50. Renner, M., Specht, C.G., and Triller, A. (2008). Molecular dynamics of postsynaptic receptors and scaffold proteins. Curr. Opin. Neurobiol. 18, 532–540. 10.1016/j.conb.2008.09.009.

51. Nakada, C., Ritchie, K., Oba, Y., Nakamura, M., Hotta, Y., Iino, R., Kasai, R.S., Yamaguchi, K., Fujiwara, T., and Kusumi, A. (2003). Accumulation of anchored proteins forms membrane diffusion barriers during neuronal polarization. Nat. Cell Biol. 5, 626–632. 10.1038/ncb1009.

52. Ewers, H., Tada, T., Petersen, J.D., Racz, B., Sheng, M., and Choquet, D. (2014). A septin-dependent diffusion barrier at dendritic spine necks. PLoS One 9, 1–19. 10.1371/journal.pone.0113916.

53. Engberg, O., Ulbricht, D., Döbel, V., Siebert, V., Frie, C., Penk, A., Lemberg, M.K., and Huster, D. (2022). Rhomboid-catalyzed intramembrane proteolysis requires hydrophobic matching with the surrounding lipid bilayer. Sci. Adv. 8, 1–10. 10.1126/sciadv.abq8303.

54. Škerle, J., Humpolíčková, J., Johnson, N., Rampírová, P., Poláchová, E., Fliegl, M., Dohnálek, J., Suchánková, A., Jakubec, D., and Strisovsky, K. (2020). Membrane Protein Dimerization in Cell-Derived Lipid Membranes Measured by FRET with MC Simulations. Biophys. J. 118, 1861–1875. 10.1016/j.bpj.2020.03.011.

55. Li, T.P., Song, Y., MacGillavry, H.D., Blanpied, T.A., and Raghavachari, S. (2016). Protein Crowding within the Postsynaptic Density Can Impede the Escape of Membrane Proteins. J. Neurosci. 36, 4276–4295. 10.1523/JNEUROSCI.3154-15.2016.

56. Liu, G., Beaton, S.E., Grieve, A.G., Evans, R., Rogers, M., Strisovsky, K., Armstrong, F.A., Freeman, M., Exley, R.M., and Tang, C.M. (2020). Bacterial rhomboid proteases mediate quality control of orphan membrane proteins. EMBO J. 39, 1–17. 10.15252/embj.2019102922.

57. Karpova, A., Hiesinger, P.R., Kuijpers, M., Albrecht, A., Kirstein, J., Andres-Alonso, M., Biermeier, A., Eickholt, B.J., Mikhaylova, M., Maglione, M., et al. (2025). Neuronal autophagy in the control of synapse function. Neuron 113, 974–990. 10.1016/j.neuron.2025.01.019.

58. Choquet, D., and Triller, A. (2013). The Dynamic Synapse. Neuron 80, 691–703. 10.1016/j.neuron.2013.10.013.

59. Gonçalves-Martins, J., Cação, C., and Ferreira, J.S. (2026). Neurexins, Ephrin receptors, and N-cadherin signaling as emerging mechanisms in synaptic dysfunction and neurodegenerative diseases. Commun. Biol. 9, 610. 10.1038/s42003-026-10023-3.

60. Hsia, H.-E., Tüshaus, J., Brummer, T., Zheng, Y., Scilabra, S.D., and Lichtenthaler, S.F. (2019). Functions of ‘A disintegrin and metalloproteases (ADAMs)’ in the mammalian nervous system. Cell. Mol. Life Sci. 76, 3055–3081. 10.1007/s00018-019-03173-7.

61. Peixoto, R.T., Kunz, P.A., Kwon, H., Mabb, A.M., Sabatini, B.L., Philpot, B.D., and Ehlers, M.D. (2012). Transsynaptic Signaling by Activity-Dependent Cleavage of Neuroligin-1. Neuron 76, 396–409. 10.1016/j.neuron.2012.07.006.

62. Kuhn, P.-H., Colombo, A.V., Schusser, B., Dreymueller, D., Wetzel, S., Schepers, U., Herber, J., Ludwig, A., Kremmer, E., Montag, D., et al. (2016). Systematic substrate identification indicates a central role for the metalloprotease ADAM10 in axon targeting and synapse function. Elife 5, 1–29. 10.7554/eLife.12748.

63. Grieve, A.G., Xu, H., Künzel, U., Bambrough, P., Sieber, B., and Freeman, M. (2017). Phosphorylation of iRhom2 at the plasma membrane controls mammalian TACE-dependent inflammatory and growth factor signalling. Elife 6, 1–22. 10.7554/eLife.23968.

64. Cavadas, M., Oikonomidi, I., Gaspar, C.J., Burbridge, E., Badenes, M., Félix, I., Bolado, A., Hu, T., Bileck, A., Gerner, C., et al. (2017). Phosphorylation of iRhom2 Controls Stimulated Proteolytic Shedding by the Metalloprotease ADAM17/TACE. Cell Rep. 21, 745–757. 10.1016/j.celrep.2017.09.074.

65. Le Gall, S.M., Maretzky, T., Issuree, P.D.A., Niu, X.-D., Reiss, K., Saftig, P., Khokha, R., Lundell, D., and Blobel, C.P. (2010). ADAM17 is regulated by a rapid and reversible mechanism that controls access to its catalytic site. J. Cell Sci. 123, 3913–3922. 10.1242/jcs.069997.

66. Scheller, J., Chalaris, A., Garbers, C., and Rose-John, S. (2011). ADAM17: a molecular switch to control inflammation and tissue regeneration. Trends Immunol. 32, 380–387. 10.1016/j.it.2011.05.005.

67. The International Mouse Phenotyping Consortium (IMPC) https://www.mousephenotype.org/.

68. Dustin, M.L., and Colman, D.R. (2002). Neural and Immunological Synaptic Relations. Science (80-.). 298, 785–789. 10.1126/science.1076386.

69. Janssen, V., and Huveneers, S. (2024). Cell–cell junctions in focus – imaging junctional architectures and dynamics at high resolution. J. Cell Sci. 137. 10.1242/jcs.262041.

70. Wilkinson, K.A., McMillan, K.J., Banks, P.J., Carmichael, R.E., Nakamura, Y., Bashir, Z.I., Cullen, P.J., and Henley, J.M. (2022). Using Lentiviral shRNA Delivery to Knock Down Proteins in Cultured Neurons and In Vivo. In Translational Research Methods in Neurodevelopmental Disorders, S. Martin and F. Laumonnier, eds. (Springer US), pp. 1–17. 10.1007/978-1-0716-2569-9_1.

71. Uno, S., Kamiya, M., Yoshihara, T., Sugawara, K., Okabe, K., Tarhan, M.C., Fujita, H., Funatsu, T., Okada, Y., Tobita, S., et al. (2014). A spontaneously blinking fluorophore based on intramolecular spirocyclization for live-cell super-resolution imaging. Nat. Chem. 6, 681–689. 10.1038/nchem.2002.

72. Künzel, U., Grieve, A.G., Meng, Y., Sieber, B., Cowley, S.A., and Freeman, M. (2018). FRMD8 promotes inflammatory and growth factor signalling by stabilising the iRhom/ADAM17 sheddase complex. Elife 7, 1–33. 10.7554/eLife.35012.

73. Bodzęta, A., Berger, F., and MacGillavry, H.D. (2022). Subsynaptic mobility of presynaptic mGluR types is differentially regulated by intra-and extracellular interactions. Mol. Biol. Cell 33, ar66. 10.1091/mbc.E21-10-0484.

74. Albrecht, D., Winterflood, C.M., and Ewers, H. (2015). Dual color single particle tracking via nanobodies. Methods Appl. Fluoresc. 3. 10.1088/2050-6120/3/2/024001.

75. Evans, A.J., Gurung, S., Wilkinson, K.A., Stephens, D.J., and Henley, J.M. (2017). Assembly, Secretory Pathway Trafficking, and Surface Delivery of Kainate Receptors Is Regulated by Neuronal Activity. Cell Rep. 19, 2613–2626. 10.1016/j.celrep.2017.06.001.

76. Mogessie, B., and Schuh, M. (2017). Actin protects mammalian eggs against chromosome segregation errors. Science 357. 10.1126/science.aal1647.

77. Martin, S., and Henley, J.M. (2004). Activity-dependent endocytic sorting of kainate receptors to recycling or degradation pathways. EMBO J. 23, 4749–4759. 10.1038/sj.emboj.7600483.

78. De Chaumont, F., Dallongeville, S., Chenouard, N., Hervé, N., Pop, S., Provoost, T., Meas-Yedid, V., Pankajakshan, P., Lecomte, T., Le Montagner, Y., et al. (2012). Icy: An open bioimage informatics platform for extended reproducible research. Nat. Methods 9, 690–696. 10.1038/nmeth.2075.

79. Schindelin, J., Arganda-Carreras, I., Frise, E., Kaynig, V., Longair, M., Pietzsch, T., Preibisch, S., Rueden, C., Saalfeld, S., Schmid, B., et al. (2012). Fiji: an open-source platform for biological-image analysis. Nat. Methods 9, 676–682. 10.1038/nmeth.2019.

80. Reese, A.L., and Kavalali, E.T. (2015). Spontaneous neurotransmission signals through store-driven Ca^2+^ transients to maintain synaptic homeostasis. Elife 4, 1–15. 10.7554/eLife.09262.001.

81. Cross, S.J., Fisher, J.D.J.R., and Jepson, M.A. (2024). ModularImageAnalysis (MIA): Assembly of modularised image and object analysis workflows in ImageJ. J. Microsc. 296, 173–183. 10.1111/jmi.13227.

82. Willems, J., de Jong, A.P.H., Scheefhals, N., Mertens, E., Catsburg, L.A.E., Poorthuis, R.B., de Winter, F., Verhaagen, J., Meye, F.J., and MacGillavry, H.D. (2020). ORANGE: A CRISPR/Cas9-based genome editing toolbox for epitope tagging of endogenous proteins in neurons. PLoS Biol. 18, e3000665. 10.1371/journal.pbio.3000665.

83. Golan, Y., and Sherman, E. (2017). Resolving mixed mechanisms of protein subdiffusion at the T cell plasma membrane. Nat. Commun. 8, 1–15. 10.1038/ncomms15851.

