## Supplementary Materials for "The unhindered lateral mobility of a rhomboid intramembrane protease enables synapse remodelling"

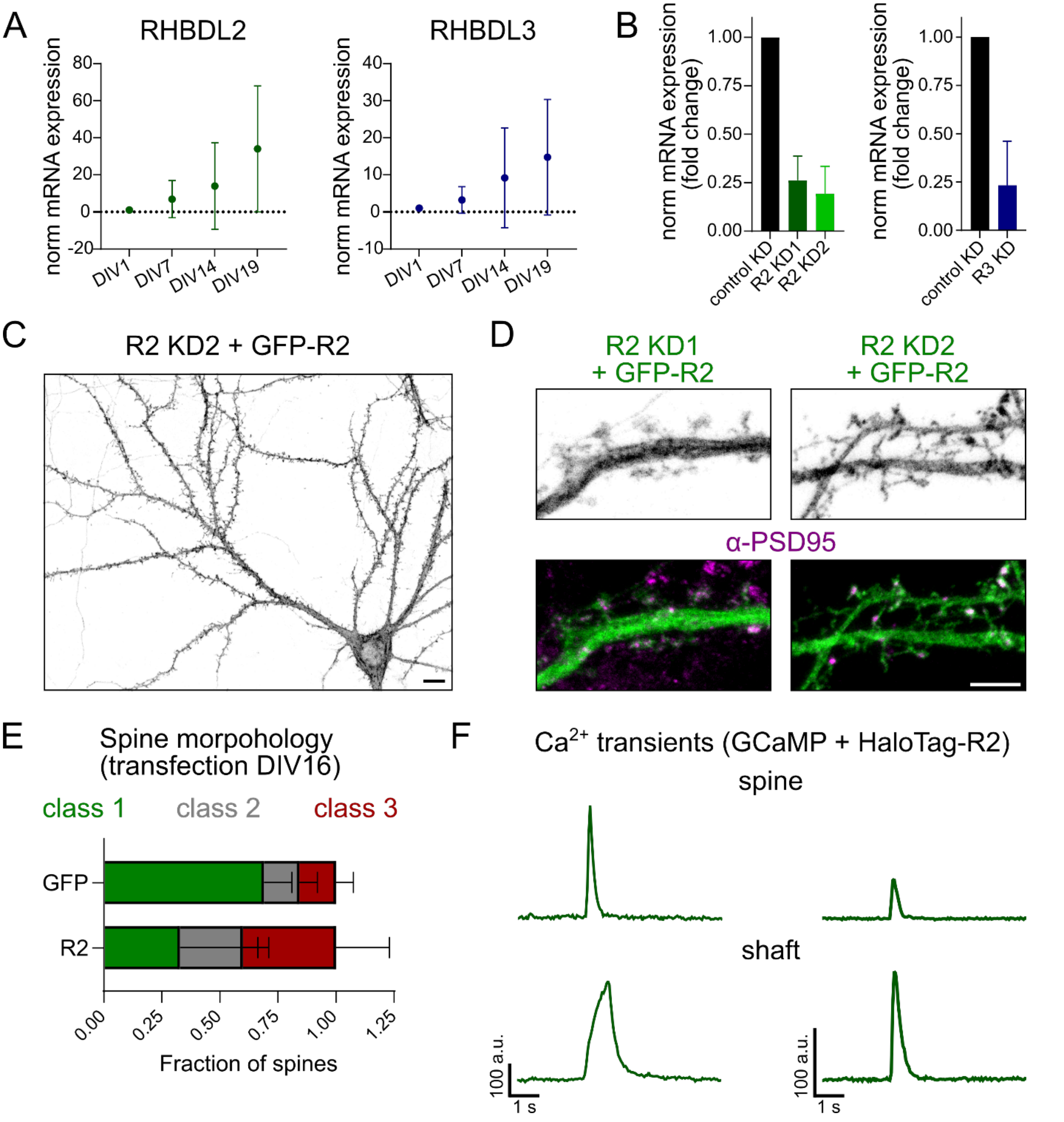


**Figure S1 (related to Figure 1).** Validation of RHBDL2 localisation and its effects on synapse function. **(A)** Graphs showing average fold change of mRNA levels of RHBDL2 and RHBDL3 in cortical neurons at different days in vitro (DIV). Changes in mRNA expression were normalised to mRNA levels at DIV1. Mean mRNA fold change ± SD (a.u.): RHBDL2: DIV 7 6.9 ± 10.5, DIV14 14.0 ± 23.4, DIV 19 34 ± 33.97; RHBDL3: DIV 7 3.2 ± 3.6, DIV14 9.2 ± 13.5, DIV 19 14.8 ± 15.6. N = 4 independent experiments. One-way ANOVA followed by Dunnett's multiple comparisons test. **(B)** Quantification of the efficiency of shRNAs used for RHBDL2 and RHBDL3 knock-down. Mean values of mRNA fold changes ± SD (a.u.): R2 KD1: 0.26 ± 0.13, R2 KD2: 0.19 ± 0.14, R3 KD: 0.23 ± 0.23. N = 2 – 6 independent experiments. **(C)** Example confocal image of a neuron with R2 KD2 and re-expression of shRNA-sensitive GFP-R2. Scale bar: 10 µm. **(D)** Example images of neurons with R2 KD1 or KD2 and re-expression of shRNA sensitive GFP-R2 co-stained with α-PSD95-S635P. Scale bar: 5 µm. Note that in **(C)** and **(D)** due to the low expression level of GFP-R2, the GFP signal was enhanced using α-GFP antibody. **(E)** Classification of spine morphology in DIV19 neurons transfected at DIV16 with GFP or GFP-R2. Details about spine classes are included in Figure 1E. Mean fraction values ± SD (a.u.): ctrl (GFP) class 1 0.69 ± 0.12, class 2 0.15 ± 0.08, class 3 0.16 ± 0.08; GFP-R2: class 1 0.33 ± 0.33, class 2 0.27 ± 0.11, class 3 0.40 ± 0.23. N = 3 independent experiments. **(F)** Examples of $F/F_0$ intensity traces of $Ca^{2+}$ transients in spines and shaft in neurons expressing HaloTag-R2.

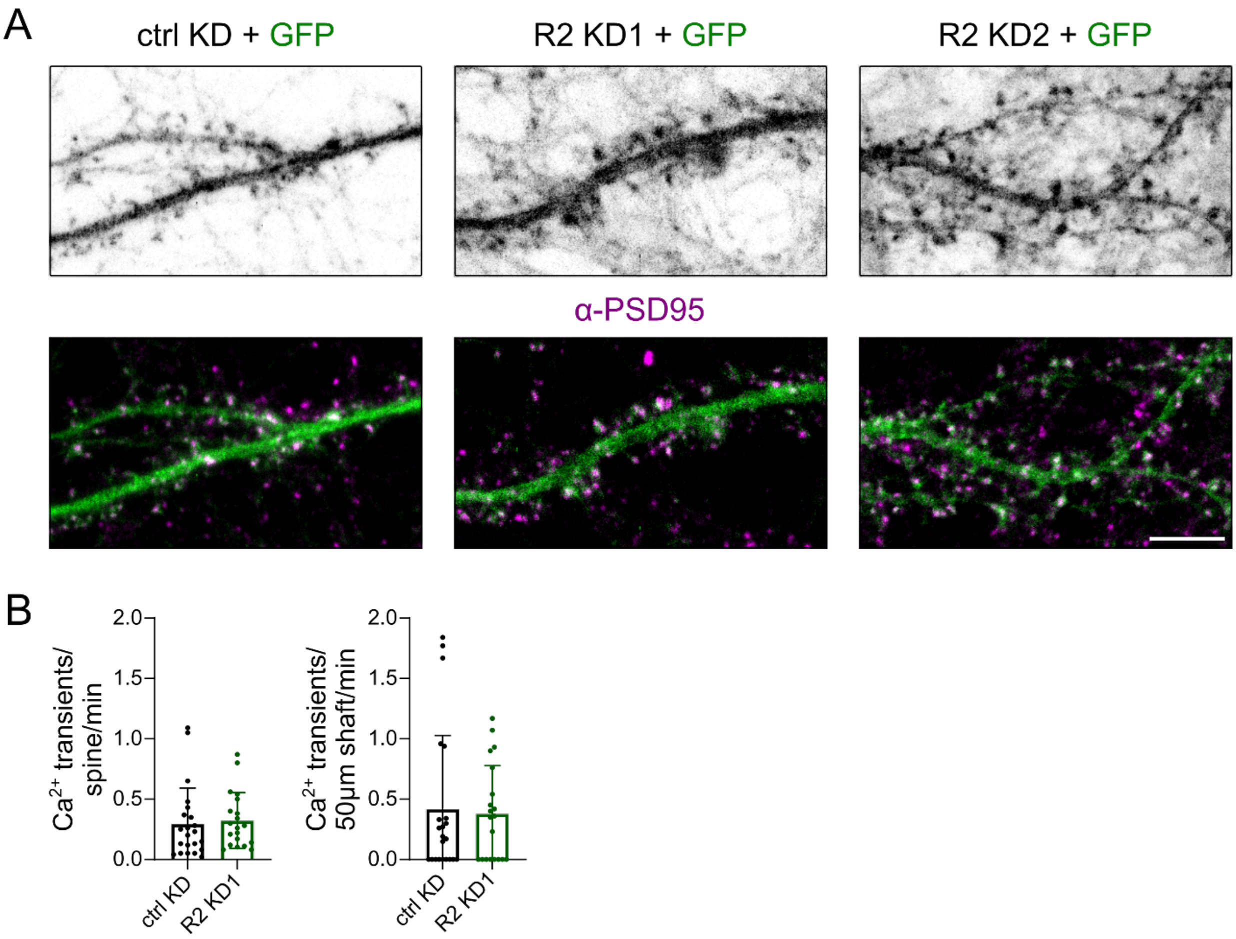


**Figure S2 (related to Figure 2).** Reduced expression of RHBDL2 does not change spine morphology or synaptic signaling. **(A)** Confocal images of neurons co-expressing GFP and control shRNA, R2 KD1 and R2 KD2 shRNAs co-stained with α-PSD95-S635P. Scale bar: 5 µm. **(B)** Quantification of GCAMP6f $Ca^{2+}$ transient frequency in neurons expressing ctrl shRNA or R2 KD1 shRNA. Mean frequency values ± SD: spine (transient/spine/min): ctrl KD 0.29 ± 0.3, R2 KD1 0.32 ± 0.23; shaft (transient/50 µm/min): ctrl KD 0.42 ± 0.61, R2 KD1 0.38 ± 0.4. N = 4 independent experiments. Unpaired t-test followed by Welch's correction.

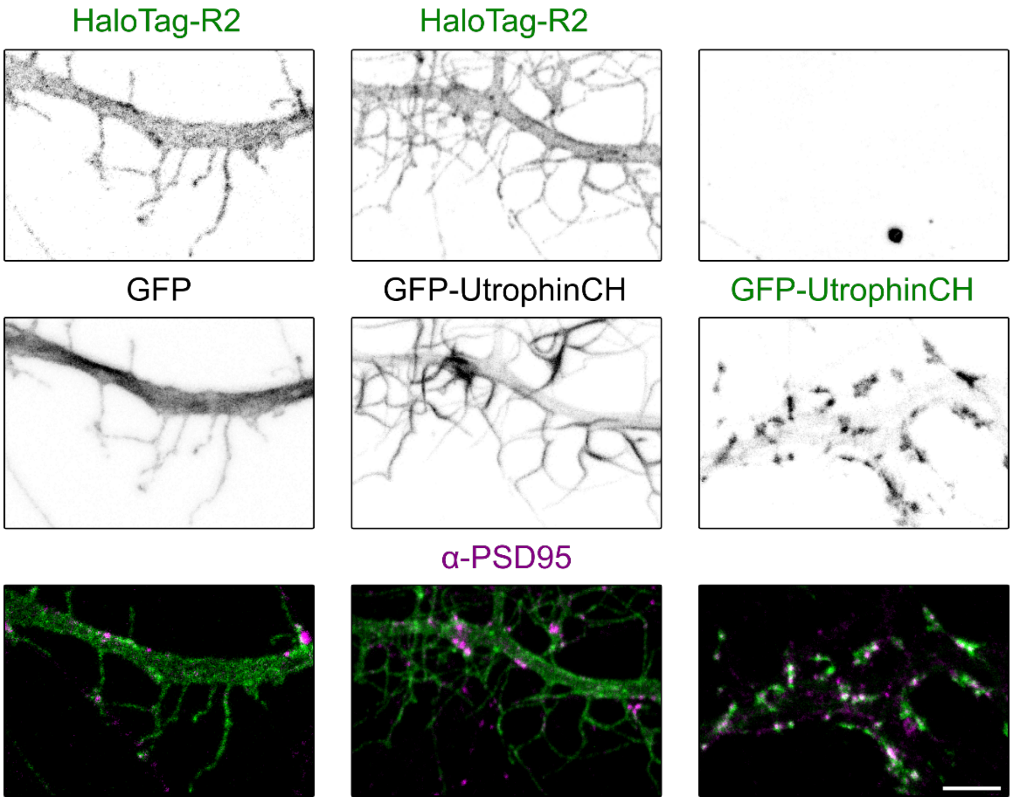


**Figure S3 (related to Figure 5).** Effect of expression of the calponin-homology domain from utrophin (UtrophinCH) on the morphology of dendritic spine. Example confocal images of neurons co-expressing HaloTag-R2 and cytosolic GFP (left), HaloTag-R2 and GFP-UtrophinCH (middle), GFP-UtrophinCH only (right) co-stained with α-PSD95-S635P. HaloTag-R2 was labeled using α-HaloTag antibody. Scale bar: 5 μm.

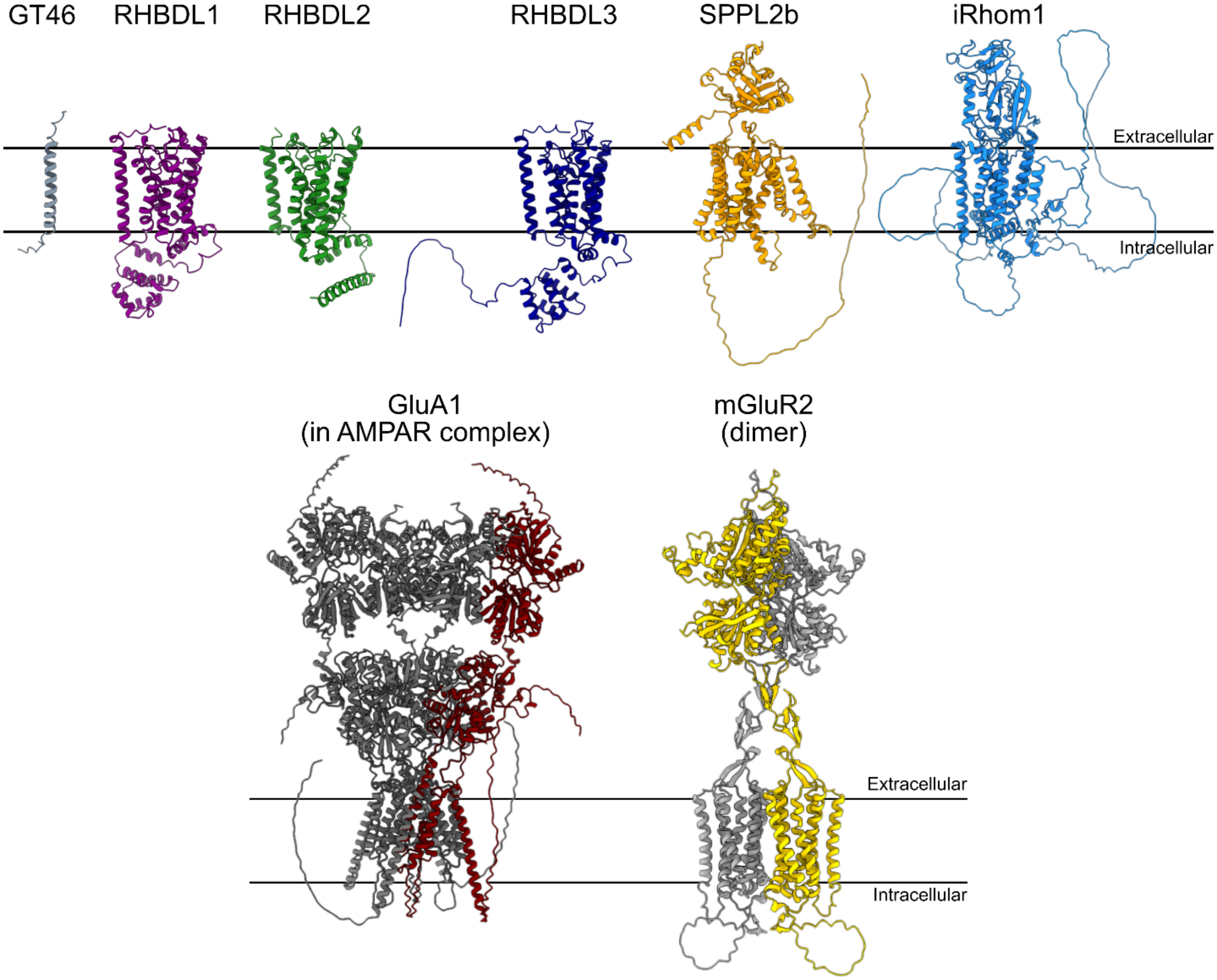


**Figure S4 (related to Figure 4).** AlphaFold2/3 predicted structural models of proteins used for SMT experiments. The model of GT46 was based on sequence encoded in plasmid HaloTag-GT46. All other models were based on rat protein sequences from UniProt: RHBDL1 – G3V8M7, RHBDL2 – D3ZA62, RHBDL3 – D4AAW6, SPPL2b – Q5PQL3, iRhom1 – Q499S9, GluA1 – P19490 (in AMPAR complex with GluA2 – P19491), mGluR2 (dimer) – P31421.

**Movie S1 (related to Figure 4)**. RHBDL2 (R2) exhibits rapid lateral diffusion at the neuronal plasma membrane. Example movie of 250 frames from of SMT time series acquired at 40Hz in stream mode HaloTag-GT46 (top) and HaloTag-R2 (bottom). Scale bar: 5 µm.

**Movie S2 (related to Figure 5).** Actin-anchored Utrophin-RHBDL2 (UR2) displayed lower mobility than wild type RHBDL2 (R2) at the dendritic plasma membrane. Example movie of 250 frames from of SMT time series acquired at 40 Hz in stream mode. HaloTag-R2 (top) and HaloTag-UR2 (bottom). Scale bar: 5 µm.
